# Maintaining oxidized H3 in heterochromatin is required for the oncogenic capacity of triple-negative breast cancer cells

**DOI:** 10.1101/2020.01.30.927038

**Authors:** Gemma Serra-Bardenys, Tian Tian, Enrique Blanco, Jessica Querol, Laura Pascual-Reguant, Beatriz Morancho, Marta Escorihuela, Sandra Segura-Bayona, Gaetano Verde, Riccardo Aiese Cigliano, Alba Millanes-Romero, Celia Jerónimo, Paolo Nuciforo, Sara Simonetti, Cristina Viaplana, Rodrigo Dienstmann, Cristina Saura, Vicente Peg, Travis Stracker, Joaquín Arribas, Josep Villanueva, Luciano Di Croce, Antonio García de Herreros, Sandra Peiró

**Author notes:** Correspondence; Vall d’Hebron Institute of Oncology, c/ Natzaret 115-117, 08035 Barcelona, Spain.

## Abstract

The histone modification of H3 oxidized at lysine 4 (H3K4ox) is catalyzed by lysyl oxidase–like 2 (LOXL2) and is enriched in heterochromatin in triple-negative breast cancer (TNBC) cells. Although H3K4ox has been linked to the maintenance of compacted chromatin, the molecular mechanism underlying this maintenance is unknown. Here we show that H3K4ox is read by the CRL4B complex, leading to the ubiquitination of histone H2A through the E3 ligase RBX1. Finally, interactions between RUVBL1/2 and LOXL2 are involved in the incorporation of the histone variant H2A.Z, which plays an essential role in the mechanism controlling the dynamics of oxidized H3. Maintenance of H3K4ox in chromatin is essential for heterochromatin properties, and disruption of any of the members involved in this pathway blocks the oncogenic properties of TNBC cells.

## INTRODUCTION

Lysyl oxidase-like 2 (LOXL2) is a histone-modifying enzyme that catalyzes the oxidation of lysine 4 in histone H3 (H3K4ox) and the generation of an aldehyde group in this amino acid position (Herranz et al., 2016). Generation of this peptidyl aldehyde can alter the local macromolecular structure as well as the nature of any protein-protein or protein-nucleic acid interactions. This is particularly relevant for gene regulation, as changes in the macromolecular status of histones can affect chromatin conformation. In fact, this oxidation is linked to transcriptional repression of several genes and heterochromatin transcripts at the onset of the epithelial-to-mesenchymal transition (EMT) and during stem cell differentiation (Herranz et al., 2016; Iturbide et al., 2015; Millanes-Romero et al., 2013). However, this modified histone is mainly enriched in heterochromatin, where it could have a role in the maintenance of condensed/compacted chromatin regions (Cebria-Costa et al., 2019). Heterochromatin is also characterized by the presence of di-methylated and tri-methylated lysine 9 in histone H3 (H3K9me2/3) and tri-methylated lysine 27 in histone H3 (H3K27me3) (Allshire and Madhani, 2018; Janssen et al., 2018). H3K9me2/3 are recognized by heterochromatin protein 1 (HP1), which has a major role in ensuring heterochromatin compaction, spreading, and inheritance (Bannister et al., 2001). Heterochromatin is not only essential for the maintenance of genome stability but can also directly affect genome integrity, through changes in its highly dynamic structure (Janssen et al., 2018). Therefore, there is a pressing need to improve our understanding of the role of heterochromatin, and the mechanisms by which it is maintained at a mechanistic level. To address this, we employed, unbiased proteomic assays to identify LOXL2 partners, as well as H3K4ox readers. We found that LOXL2 interacts with members of the SRCAP and TIP60 complexes, and that H3K4ox is read by the CRL4B complex. Both of these complexes are involved in loading the histone H2A variant, H2A.Z, into chromatin (Buschbeck and Hake, 2017; Morrison and Shen, 2009). We also discovered that the CRL4B complex is involved in the ubiquitination of H2A by RBX1, that is required for the exchange of the H2A variant, H2A.Z. Notably, both *in vitro* and *in vivo*, these series of events are necessary to maintain condensed chromatin in triple-negative breast cancer (TNBC) cells. Knocking down any component of this pathway resulted in inhibition of oncogenic and metastatic traits in TNBC cells, including proliferation, migration, invasion, and *in vivo* tumor formation capacity. Thus, we propose the existence of a direct link between oxidation of H3, chromatin condensation, and tumorigenic capacities, and argue that any of the components described in the H3K4ox pathway could potentially be targetable factors for treating TNBC.

## RESULTS

### The Roles of LOXL2, RUVBL2, and H2A.Z in Chromatin Condensation

We revisited a tandem-affinity purification approach previously published by our lab to identify putative LOXL2 interactors that might be involved in the maintenance or generation of compacted chromatin induced by H3K4ox-LOXL2 (Cebria-Costa et al., 2019; Herranz et al., 2016). Analysis of the new list of interactors showed the presence of two AAA+ ATPases (RUVBL1 and RUVBL2), DMAP1 and BAF53, all of which are members of the SRCAP and TIP60 complexes (Figure 1A and Table S1). We confirmed their interactions with the complex by co-immunoprecipitation experiments. Specifically, ectopically-expressed LOXL2-FLAG and endogenous RUVBL1, RUVBL2, BAF53, and DMAP1 interacted in HEK293T cells (Figure 1B). RUVBL1/2 proteins are essential for the proper assembly and functionality of the chromatin remodeler complex (Cheung et al., 2010; Jonsson et al., 2004; Nguyen et al., 2013). Notably, the presence of RUVBL2 (but not RUVBL1) in the chromatin remodeler complex is required to induce conformational changes that allow ADP-ATP exchange (Cheung et al., 2010). We therefore performed lost-and gain-of-function experiments in the presence or absence of RUVBL2 to determine whether this complex is important in H3K4ox-mediated chromatin condensation. HEK293T cells infected with a lentivirus carrying specific shRNA for RUVBL2 (shRUVBL2) revealed that the absence of RUVBL2 led to the loss of the LOXL2 interaction (Figure 1C). As previously observed (Izumi et al., 2010; Morwitzer et al., 2019; Venteicher et al., 2008), the levels of the RUVBL1 protein, but not of its RNA, decreased in the absence of RUVBL2 (Figures 1C and S1A).

**Figure 1.**
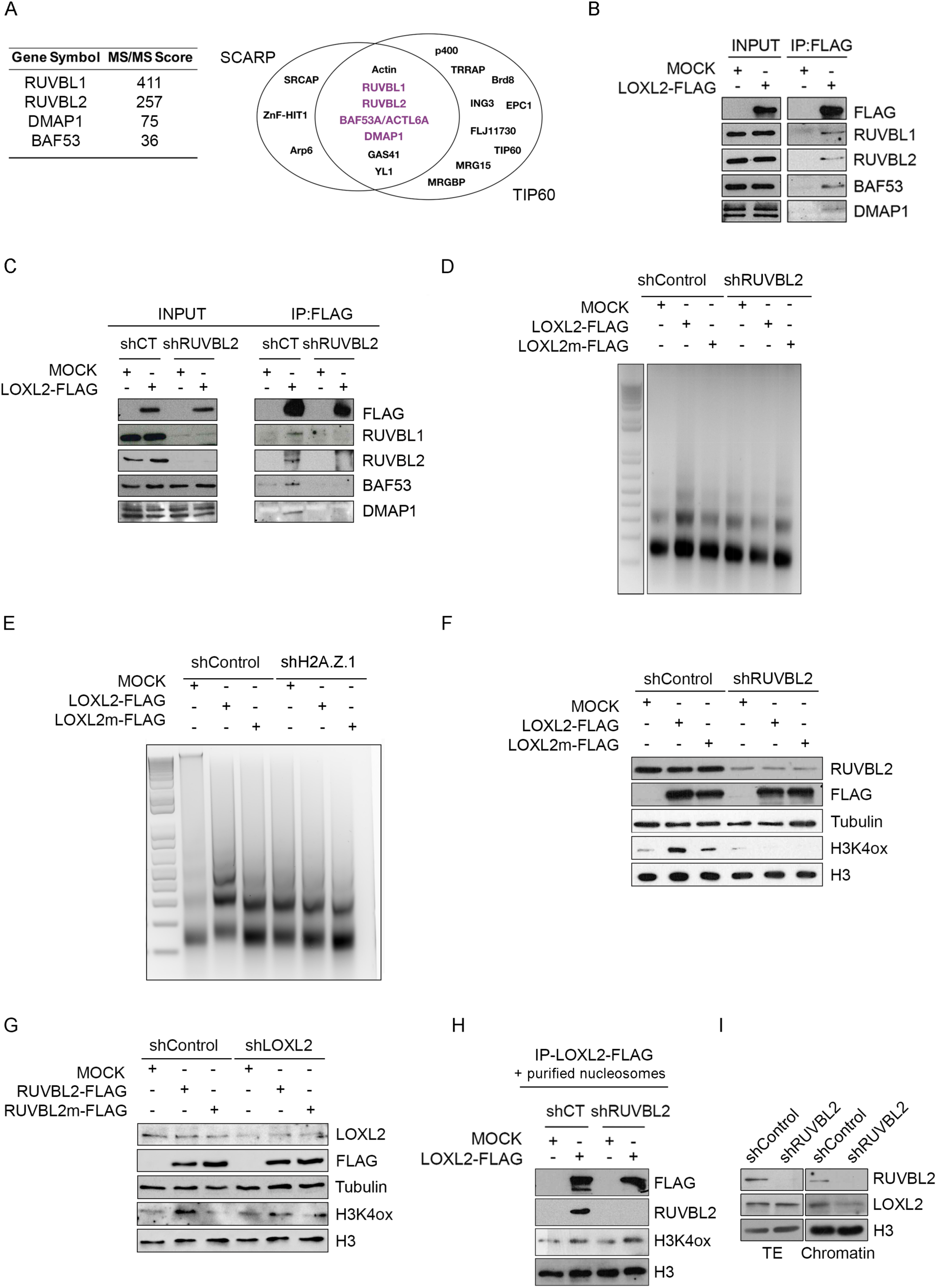
LOXL2 interacts with members of the SRCAP and TIP60 complexes, and RUVBL2 is required for H3K4ox maintenance and the induction of chromatin compaction. **A)** Putative LOXL2 interactors identified in a previously-published tandem-affinity purification approach and mass spectrometry (MS) analysis. Left panel, MS scores; right panel, schematic representation of SRCAP and TIP60 complexes with the identified LOXL2 partners (in purple). **B, C)** HEK293T cells either not infected (**B**) or infected with shControl or shRUVBL2 (**C**) were transiently transfected with LOXL2-Flag or an empty vector, and 48 hr post-transfection extracts were immunoprecipitated (IP) with anti-Flag-M2 beads. Immunocomplexes were analyzed by Western blot with the indicated antibodies. **D, E)** HEK293T cells infected with shControl, shRUVBL2 (**D**), or shH2A.Z.1 (**E**) were transfected with wild-type LOXL2 (LOXL2), an inactive LOXL2 mutant (LOXL2m), or an empty vector (mock). At 48 hr post-transfection, isolated nuclei were digested with micrococcal nuclease (MNase) for 2 min, and total genomic DNA was analyzed by agarose gel electrophoresis. **F)** HEK293T cells infected with shControl or shRUVBL2 were transfected with wild-type LOXL2 (LOXL2), inactive mutant LOXL2 (LOXL2m), or an empty vector (mock). At 48 hr post-transfection, total and histone extracts were analyzed by Western blot with the indicated antibodies. **G)** HEK293T cells infected with shControl or shLOXL2 were transfected with wild-type RUVBL2 (RUVBL2), an ATPase-deficient RUVBL2 mutant (RUVBL2m), or an empty vector (mock). At 48 hr post-transfection, total and histone extracts were analyzed by Western blot with the indicated antibodies. **H)** HEK293T cells infected with shControl or shRUVBL2 were transfected with an empty vector or LOXL2-FLAG. At 48 hr post-transfection, co-immunoprecipitation assays with anti-Flag-M2 beads were performed. Immunoprecipitated complexes were then incubated with purified nucleosomes for 2 hr at 37°C, and samples were analyzed by Western blot with the indicated antibodies. **I)** MDA-MB-231 cells were infected with shControl or shRUVBL2. Total extracts and chromatin fraction obtained by subcellular fractionation assays were analyzed by Western blot with the indicated antibodies. Note that in Figures 1B, 1C, 1D, and 1I, the intervening lanes were removed as indicated.

To determine the effects on chromatin condensation, we next analyzed the general chromatin status of cells with or without active LOXL2. MNase digestion experiments in HEK293T cells transfected with wild-type LOXL2 showed increased global chromatin compaction as compared to cells transfected with a mutant LOXL2 (LOXL2m) with compromised catalytic activity (Herranz et al., 2016) (Figure S1B). Notably, this increase in chromatin compaction upon overexpressed LOXL2 only occurred in the presence of RUVBL2 but not after RUVBL2 had been knocked-down (using shRUVBL2) (Figure 1D), suggesting that RUVBL2 and the maintenance of the complex are key to the LOXL2-mediated formation of compacted chromatin. One of the main functions of the SRCAP and TIP60 complexes is the incorporation of H2A.Z into chromatin. In vertebrates, two isoforms of H2A.Z exist, H2A.Z.1 and H2A.Z. 2 coded by two different genes (*H2AFZ* and *H2AFV*). Most of the studies do not distinguish between the two isoforms, therefore what is commonly referred as H2A.Z is the H2A.Z.1 isoform. H2A.Z is a histone variant that is enriched in heterochromatin, and therefore present in compacted chromatin regions. In agreement with this, knocking down H2A.Z (Figure S1C) blocked chromatin compaction that was induced by LOXL2 overexpression (Figure 1E), suggesting that RUVBL2 and H2A.Z exchange play a role in LOXL2-induced chromatin condensation.

It is possible that RUVBL2 and H2A.Z are necessary for the formation of H3K4ox by LOXL2 or for LOXL2 chromatin localization (and thus have indirect rather than direct roles in the maintenance of compacted chromatin). To test this hypothesis, we performed a series of *in vivo* and *in vitro* biochemical assays in HEK293T cells. We first observed that the absence of RUVBL2 or H2A.Z (by shRNA depletion) blocked transfected LOXL2 from oxidizing H3 (Figures 1F and S1D). To elucidate whether oxidation required the catalytic activity of RUVBL2, shLOXL2 cells (or shControl cells) were transfected with wild-type RUVBL2 or a mutant RUVBL2 with no ATPase activity (RUVBL2m) (Grigoletto et al., 2013; Puri et al., 2007). Western blot analysis of H3K4ox levels revealed that H3 was oxidized only when both LOXL2 and wild-type RUVBL2 (but not RUVBL2m) were present (Figure 1G). We have previously shown that *in vitro* reactions with recombinant LOXL2 and purified histones or nucleosomes do not require RUVBL2 for catalyzing H3K4ox (Herranz et al., 2016). Thus, we speculated that RUVBL2 is needed for maintaining oxidized H3 in chromatin. For instance, by regulating H3 exchange and/or for bringing LOXL2 physically to chromatin, rather than for the LOXL2 enzymatic activity. Indeed, *in vitro* reactions with the immunoprecipitated LOXL2 complex (from shControl or shRUVBL2 cells) and purified nucleosomes showed enzymatic activity of the LOXL2 complex in oxidizing nucleosomes in the absence of RUVBL2 (Figure 1H).

To determine whether RUVBL2 is necessary to recruit and maintain LOXL2 to chromatin, we analyzed the levels of LOXL2 and H3K4ox in the TNBC cell line MDA-MB-231 by subcellular fractionation; note that this breast cancer cell line was selected because it expresses high levels of LOXL2 and exhibits high H3K4ox (Cebria-Costa et al., 2019). Indeed, in shRUVBL2 cells, reduced levels of LOXL2 were detected in the chromatin fraction (as compared to shControl cells), suggesting that the presence of RUVBL2 is important to maintain LOXL2 into chromatin (Figure 1I). Further, no changes in LOXL2 chromatin levels were observed in the absence of H2A.Z, suggesting that H2A.Z is not involved in either the recruitment or maintenance of LOXL2 at chromatin (Figure S1E).

We next analyzed whether RUVBL2 is involved in the dynamism of H3. To that end, we monitored the deposition of newly synthesized histones H3.1 and H3.3 using the SNAP-tag system (Bodor et al., 2012; Jansen et al., 2007; Ray-Gallet et al., 2011). After the quench-chase-pulse assay (Figures 2A and S2), MDA-MB-231 cells stably expressing H3.1-SNAP or H3.3-SNAP were treated with Triton X-100 before fixation, to visualize only chromatin-incorporated histones. We observed a decrease both in H3.1 and H3.3 deposition in the absence of either RUVBL2 or LOXL2, indicating that (i) when the levels of H3K4ox are high, there is a constant exchange of the H3 variants, and that (ii) the presence of oxidized H3 signals this exchange (Figures 2A and 2B). In summary, we found that LOXL2 interacts with core members of the SRCAP and TIP60 complexes, and that active RUVBL2 and H2A.Z incorporation are essential to induce LOXL2-mediated chromatin condensation and to maintain H3K4ox levels into chromatin.

**Figure 2.**
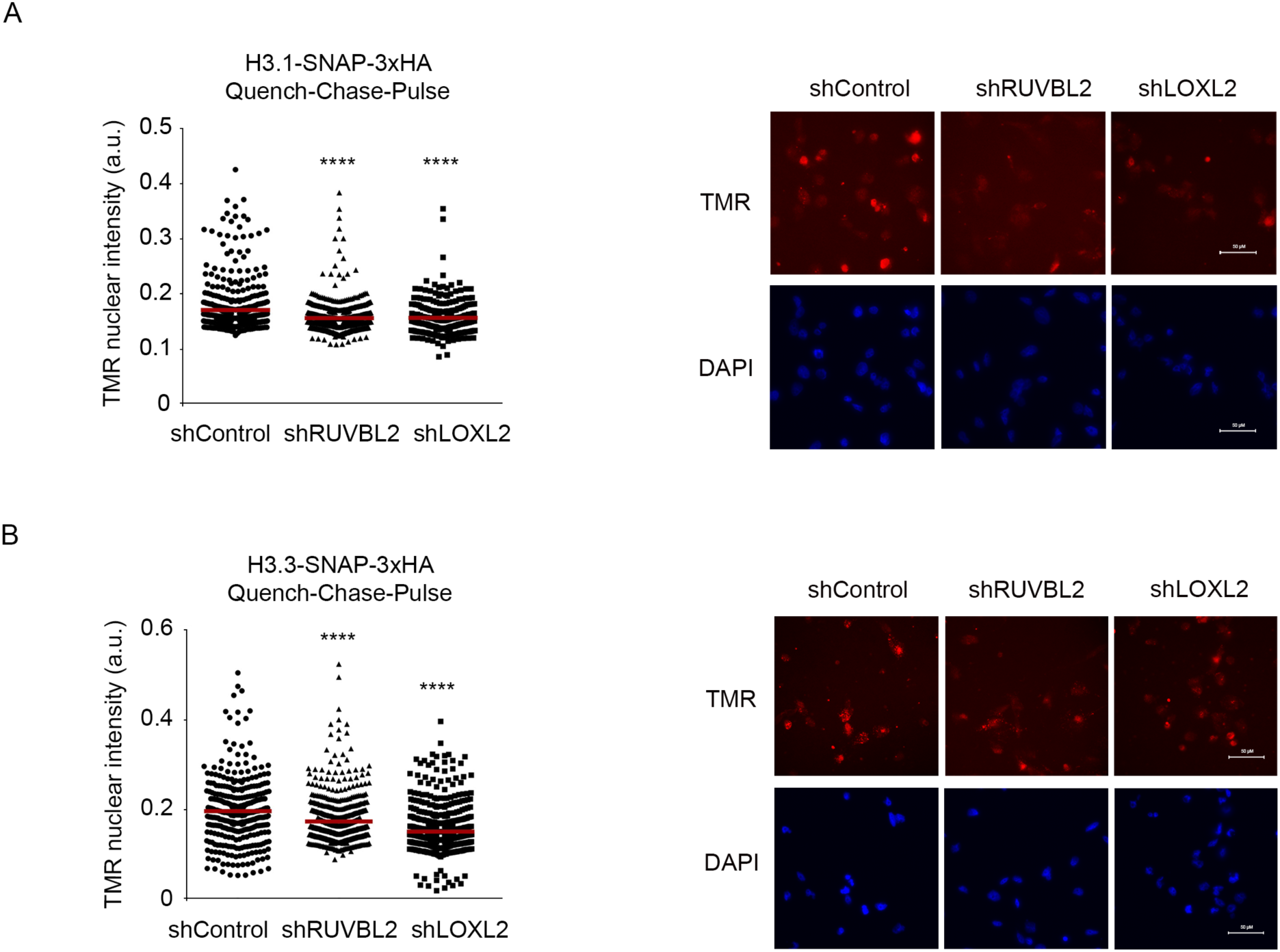
RUVBL2 or LOXL2 knockdown reduces the H3.1 and H3.3 exchange. MDA-MB-231 cells stably expressing H3.1-SNAP **(A)** or H3.3-SNAP **(B)** were infected with shControl, shLOXL2, or shRUVBL2. After 96 h, quench-chase-pulse experiments were performed to specifically label newly-synthetized histones. Cells were quenched with 5 µM SNAP-cell Block for 30 min at 37°C, washed, and incubated in fresh medium at 37°C for the chase period (7 h). The pulse step was performed with 2 µM SNAP-cell TMR Star for 15 min. Cells were pre-extracted in 0.2% Triton X-100 to remove unbound chromatin fraction and fixed with paraformaldehyde. Nuclei were stained with DAPI. High-throughput microscopy (HTM)-mediated quantification of the nuclear intensity of SNAP-H3.1/3 was performed. Quantification and images of one representative experiment are shown (at least 130 nuclei were analyzed). At least two independent experiments were performed. Scale bars represent 50 µM. Median is indicated in red. *** p < 0.001. **** p < 0.0001.

### H3K4ox, RUVBL2, and H2A.Z are Enriched in Constitutive Heterochromatin

We next used ChIP-seq to determine if H3K4ox, RUVBL2, and H2A.Z are located in the same chromatin regions (in MDA-MB-231 cells) (note that commercially-available LOXL2-specific antibodies do not work for either ChIP-seq or ChIP-PCR). As H3K4ox is a histone modification enriched in heterochromatin (Cebria-Costa et al., 2019), multi-locus reads were included during the ChIP-seq mapping. The analysis revealed strong overlap between H3K4ox and RUVBL2 (38% and 60% of total H3K4ox and RUVBL2 peaks, respectively). Although the overlap between H3K4ox and RUVBL2 common regions and H2A.Z was modest (26% of total H2A.Z peaks) we found up to 10213 genomic regions presenting H3K4ox, RUVBL2 and H2AZ co-occupancy (Figure 3A,3B and 3C). Gene ontology (GO) analysis of the 2267 genes located within the 10213 regions revealed a significant enrichment in regulation of calcium signaling, leukocyte differentiation, or nucleotide catabolism (Figure 3D). Transcriptome analysis by RNA-seq revealed that 30% of genes dysregulated in shLOXL2 were also differentially expressed in shRUVBL2 (Figure 3E). The expression changes observed in a selection of these genes were further confirmed by RT-PCR (Figures S3C and S3D). GO analysis showed that differentially expressed genes were mostly enriched in pathways controlling DNA replication and cell proliferation (Figures S3A and S3B). Notably, we observed that these genes were not located in chromatin regions enriched with H3K4ox and RUVBL2, suggesting that H3K4ox and RUVBL2 were not implicated directly in gene expression regulation but rather in chromatin structure (Figure 3F). Validation of ChIP-seq was performed by conventional ChIP-PCR in shLOXL2 and shRUVBL2 for selected genomic regions. Importantly, we observed that, in the absence of LOXL2 and RUVBL2, both H3K4ox and H2A.Z levels decreased (Figures 4A,4B and S4), suggesting that the presence of H3K4ox is essential to maintain H2A.Z in chromatin.

**Figure 3.**
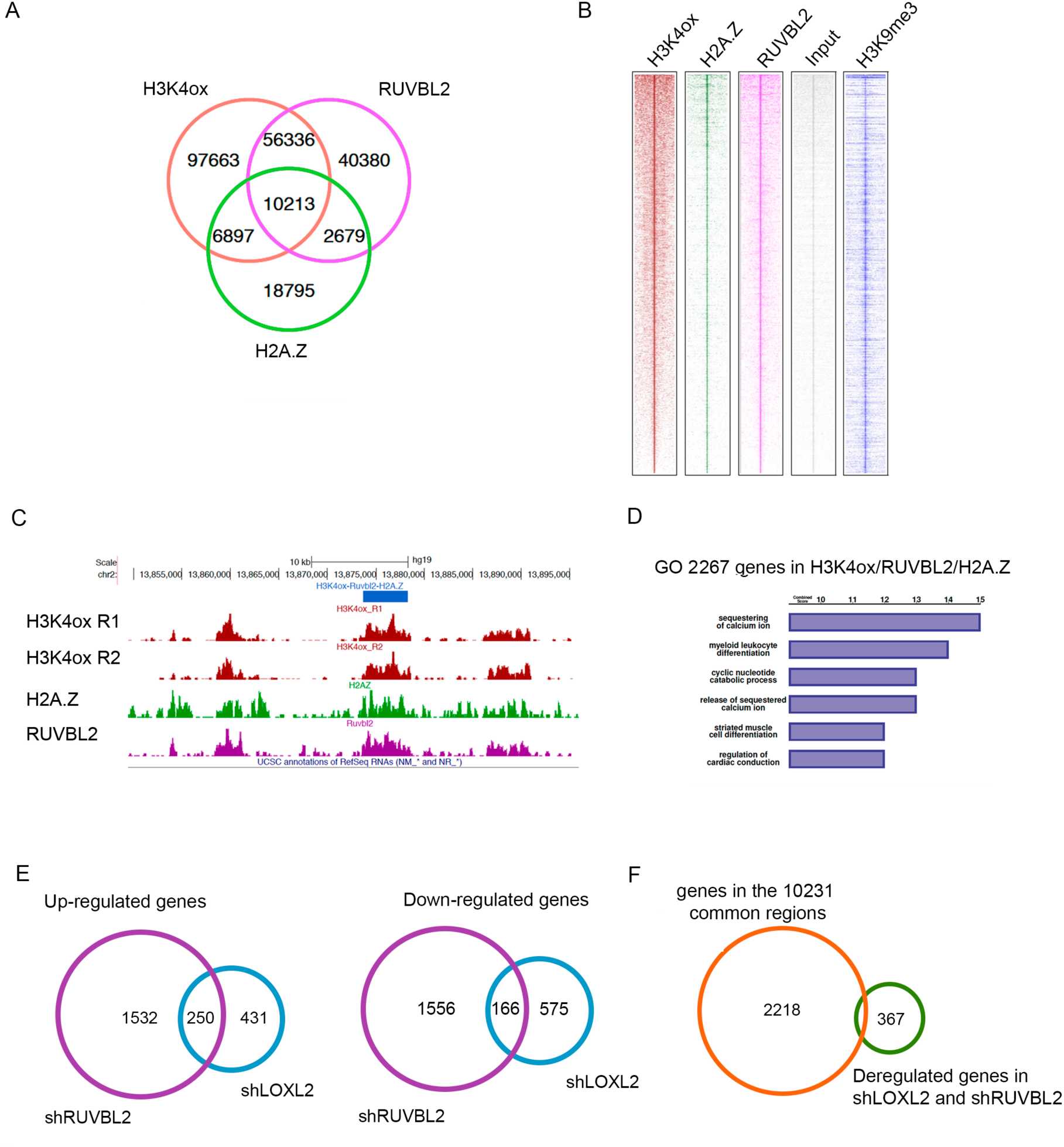
H3K4ox, RUVBL2, and H2A.Z are enriched in constitutive heterochromatin but are not directly implicated in gene expression regulation. **A)** Venn diagram showing the overlap between H2A.Z, RUVBL2, and H3K4ox ChIP-sequencing peaks. **B)** Heatmap of H3K4ox, H2A.Z, RUVBL2, input, and H3K9me3 (ChIP from GSE85158) in the 10,231 common genomic regions. **C)** UCSC Genome Browser overview of one region in chromosome 2 containing ChIP-seq profiles of H3K4ox (2 replicates, in red), H2A.Z (in green), and RUVBL2 (in purple). **D)** GO analyses (Biological Process 2018) of 2,267 genes located in the common regions. **E)** Venn diagram showing the overlap between upregulated (left) and downregulated (right) genes in MDA-MB-231 cells knocked down for RUVBL2 (shRUVBL2) or for LOXL2 (shLOXL2). **F)** Venn diagram showing the overlap between the 2,267 genes embedded in the 10,231 genomic regions with H3K4ox, H2A.Z, and RUVBL2 (orange) and the common genes up- or downregulated in MDA-MB-231 cells knocked down for RUVBL2 (shRUVBL2) and LOXL2 (shLOXL2) (green).

**Figure 4.**
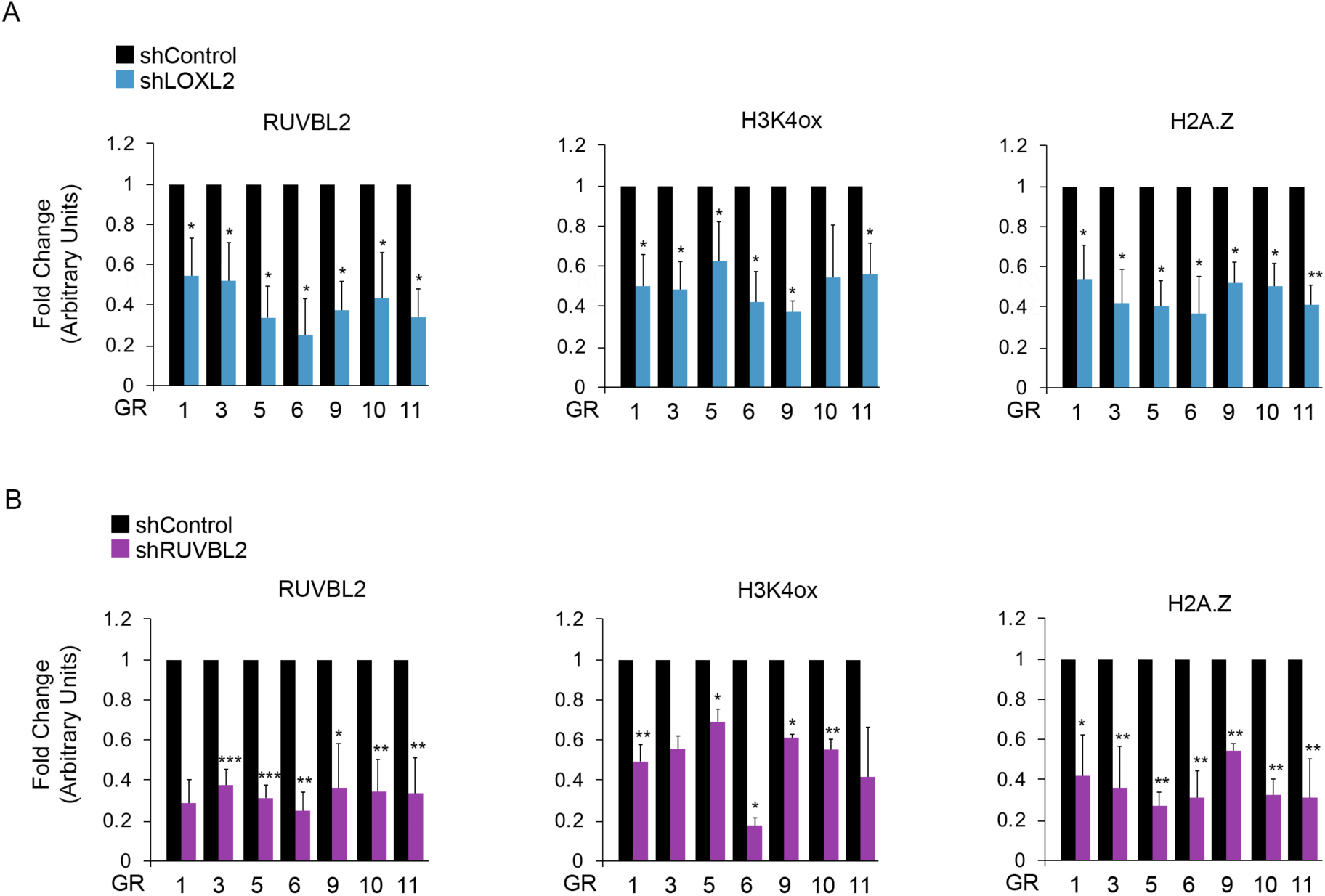
Incorporation of H2A.Z and H3K4ox into heterochromatin regions depend on LOXL2 and RUVBL2. MDA-MB-231 cells were infected with shControl, shLOXL2 **(A**), or shRUVBL2 **(B)**. ChIP-PCR experiments of RUVBL2, H3K4ox, and H2A.Z were performed in selected genomic regions (GR). Data of qPCR amplifications were normalized to the input and to total immunoprecipitated H3 for H3K4ox. Results are expressed as a fold-change relative to the data obtained in shControl, which was set to 1. Error bars indicate SD in at least three experiments. *p < 0.05, **p ≤ 0.01, ***p ≤ 0.001.

### DDB1 is a H3K4ox Reader Involved in the Ubiquitination of Histone H2A Through the CRL4B Complex

To decipher the molecular mechanism(s) by which LOXL2-mediated H3K4ox works together with RUVBL2 and H2A.Z to maintain chromatin condensed, we first identified H3K4ox readers by mass spectrometry (MS)–based proteomics. As the aldehyde group generated in the histone H3 tail after the LOXL2 reaction on trimethylated lysine 4 is highly reactive (Herranz et al., 2016), it is not possible to produce an H3 peptide with this modification. Therefore, we used a biotinylated-H3 peptide containing an intermediate alcohol in position 4 (using the artificial amino acid 6-hydroxynorleucine) that resembles the intermediate product oxidized during the LOXL2 reaction. This peptide was used for pull-down experiments (Figure 5A). As a control, we used the same amount of an immobilized biotinylated-irrelevant peptide (Figures S5A and S5B). Proteins from MDA-MB-231 nuclear extracts that bound immobilized peptides were digested with trypsin, and the resulting peptides were analyzed by liquid chromatography-tandem mass spectrometry (LC-MS/MS) (Figure S5A). We identified 77 putative H3K4ox readers in MDA-MB-231 nuclear extracts from the peptide-pulldown interaction plot, with WD-repeat (WDR)-domain–containing proteins the most enriched (Figure 5B and Table S2). Additionally, we identified the DNA damage–binding protein 1 (DDB1), a component of Cullin4-containing E3-ubiquitin ligase complexes (CRL4s). In these complexes, DDB1 binds to CUL4A or CUL4B to recruit CUL4-DDB1 associated factors (DCAF), a family of WDR-domain containing proteins that confer substrate specificity to the complex (Angers et al., 2006; He et al., 2006; Higa et al., 2006). DDB1 also binds to a RING finger protein (RING1B or RBX1) and enables interaction with the E2-conjugating enzyme. A large number of CRL4 complexes have been identified so far, exhibiting specificity for H3, H4 and/or H2A in many cases (Jackson and Xiong, 2009; Wang et al., 2006). Two complexes that contain DDB1 have been described to monoubiquitinate histone H2A (producing H2Aub): the DDB1-CUL4B-RBX1 (CRL4B) complex and the DDB1-DDB2-CUL4A-CUL4B-RING1B (UV-RING1B) complex (Gracheva et al., 2016; Hu et al., 2012; Liu et al., 2018). H2Aub mediates transcription repression and regulates chromatin accessibility during DNA repair in response to UV-induced DNA damage. The CRL4B complex is also involved in transcription repression through its monoubiquination of K119 in H2A (H2AK119ub) (Hu et al., 2012). *In vitro* experiments revealed that DDB1 directly binds H3K4ox, thus establishing its activity as a reader (Figure 5C). Moreover, DDB1 was recruited into the chromatin fraction when LOXL2 was overexpressed (Figure S5C, left panel). LOXL2 knock-down and the concomitant decrease in H3K4ox levels in the chromatin fraction also caused a decrease in DDB1 levels, further supporting its role as a H3K4ox reader (Figure S5C, right panel). Co-immunoprecipitation experiments from MDA-MB-231 cell extracts showed that DDB1 binds H3K4ox even in the presence of ethidium bromide (EtBr), revealing that this interaction is independent of DNA (Figure 5D). Reverse-IP pull-downs showed that DDB1 interacted with H3K4ox, RING1B, CUL4B and RBX1 (Figure 5E, upper panel), and that H3K4ox interacted with RING1B, CUL4B, RBX1 and DDB1 (Figure 5E, lower panel).

**Figure 5.**
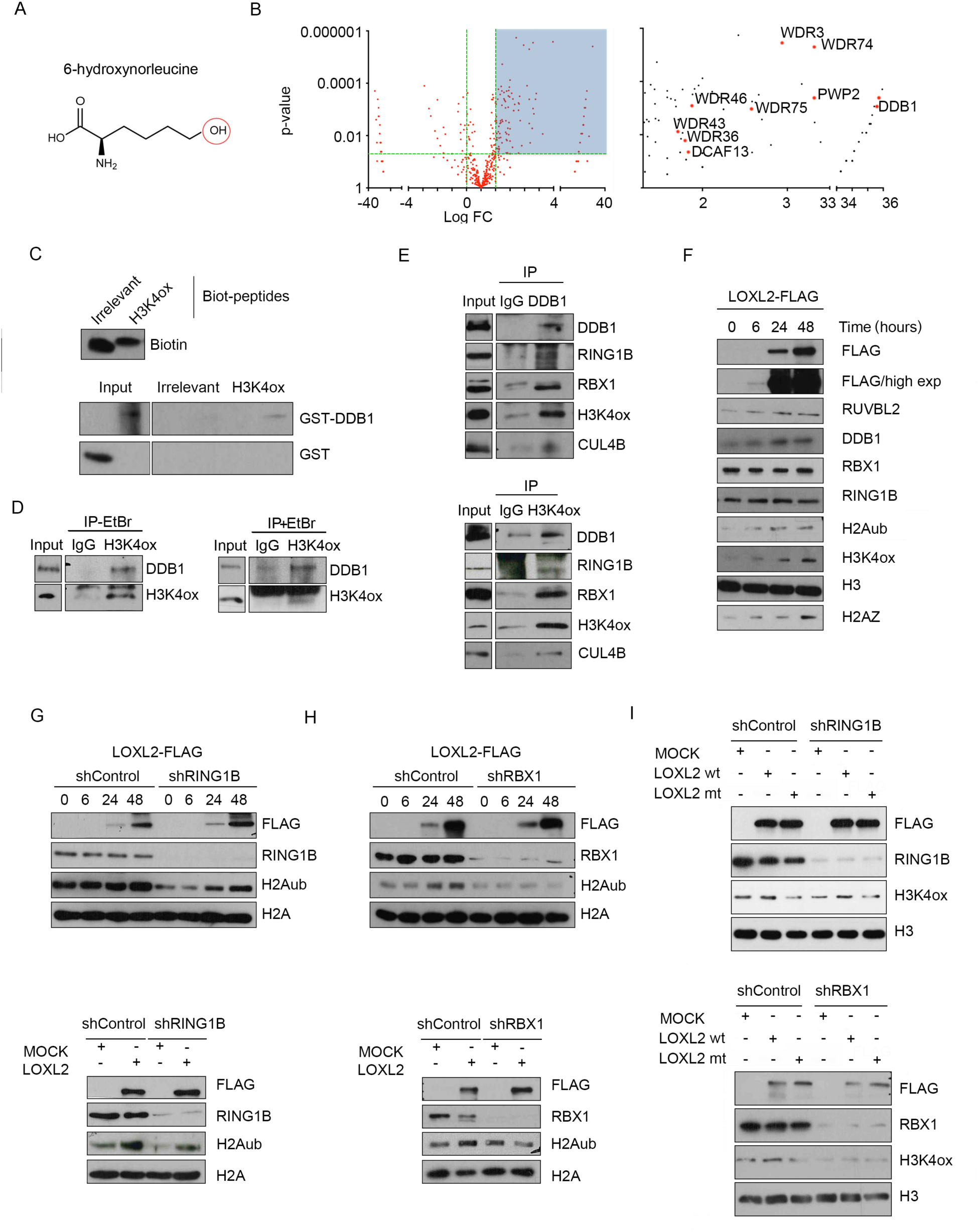
DDB1 is an H3K4ox reader involved in the ubiquitination of histone H2A through the CRL4B complex. **A)** The artificial amino acid 6-hydroxy-norleucine, which resembles the intermediate alcohol that is oxidized during LOXL2 reaction, was added to position 4 of the biotinylated H3 peptide. **B)** Peptide pull-down interaction plot, showing the putative H3K4ox readers in the upper right part (right panel). The zoom of the highlighted graph area shows the identified proteins (left panel). Background proteins are clustered together around the origin of the plot. **C)** H3K4ox or irrelevant peptides were immobilized onto magnetic streptavidin beads and incubated with DDB1-GST or GST. Complexes were washed, and proteins bound to streptavidin beads were analyzed by Western blot. In parallel, recovery of peptides was analyzed by Tricine-SDS-PAGE and incubation with streptavidin-HRP. **D)** Nuclear extracts from MDA-MB-231 cells were immunoprecipitated with anti-H3K4ox in the presence or absence of ethidium bromide. Immunocomplexes were analyzed by Western blot with the indicated antibodies. **E)** Nuclear extracts from MDA-MB-231 cells were immunoprecipitated with anti-DDB1 (upper panel) or anti-H3K4ox (lower panel). Immunocomplexes were analyzed by Western blot with the indicated antibodies. **F)** HEK293T cells were transfected with LOXL2-FLAG. Chromatin association assays were performed at 0, 6, 24, and 48 hr post-transfection, and indicated proteins were analyzed by Western blot with specific antibodies. **G, H)** HEK293T cells were infected with shControl, shRING1B (**G**), or shRBX1 (**H**). After selection, cells were transfected with LOXL2-FLAG, and chromatin association assays were performed at 0, 6, 24, and 48 hr post-transfection (upper panels). Additionally, infected cells were transfected with LOXL2-FLAG or an empty vector to do subcellular fractionation assays after 48 hr (lower panels). Protein levels in the different fractions were analyzed by Western blot with the indicated antibodies. **I)** HEK293T cells infected with shControl, shRING1B (upper panel) or shRBX1 (lower panel) were transfected with wild-type LOXL2 (LOXL2), an inactive LOXL2 mutant (LOXL2m), or empty vector (mock). At 48 hr post-transfection, histone and total extracts were analyzed by Western blot with the indicated antibodies. Note that, in Figures 5C, 5D, and 5E, the intervening lanes were removed as indicated.

To assess if DDB1 recruitment to chromatin depends on LOXL2 and H3K4ox, LOXL2 was transfected in HEK293T cells, and the chromatin fraction was obtained at different time points after transfection. In parallel with LOXL2 expression after transfection, we observed an increase in H3K4ox, the concomitant recruitment of RUVBL2, DDB1 and H2A.Z to chromatin, and increased levels of H2Aub (Figure 5F). In contrast, RING1B and RBX1 levels remained constant in chromatin throughout the experiment (Figure 5F). Subcellular fractionation assays also showed an increase in ubH2A levels when LOXL2 is overexpressed (Figure S5D). To determine whether LOXL2-dependent ubiquitination of H2A relies on RINGB and/or RBX1, chromatin assembly and subcellular fractionation experiments were performed using shRING1B or shRBX1 (MDA-MB-231) cells. LOXL2-dependent H2Aub occurred even in the absence of RING1B (Figure 5G) but was completely abrogated in the absence of RBX1 (Figure 5H). Accordingly, maintenance of H3K4ox levels after LOXL2 overexpression required RBX1 but not RING1B (Figure 5I). Finally, Western blot showed that ectopic expression of RBX1 rescued H2Aub levels in the presence of LOXL2 in shRBX1 cells, confirming its requirement for this modification (Figure S5E).

### H2Aub is Required to Maintain Compacted Chromatin

Is the presence of H2Aub required for maintaining the H2A.Z levels in heterochromatin? As expected, H2Aub levels of selected genomic regions decreased in the absence of LOXL2 or RBX1, with a concomitant decrease in H2A.Z occupancy in the same regions (Figure 6A, 6B and S4). Notably, the amount of RUVBL2 recruited to chromatin did not change in the absence of RBX1, suggesting that the decreased occupancy of H2A.Z was due to the reduced levels of H2Aub (Figure 6A and 6B). General chromatin compaction analyzed by MNase assay showed that, in the absence of RBX1, although LOXL2 was still recruited to chromatin did not induce chromatin compaction (in agreement with the observation that reduced levels of RBX1 caused reduced levels of H3K4ox in chromatin) (Figures 6C and S5F).

**Figure 6.**
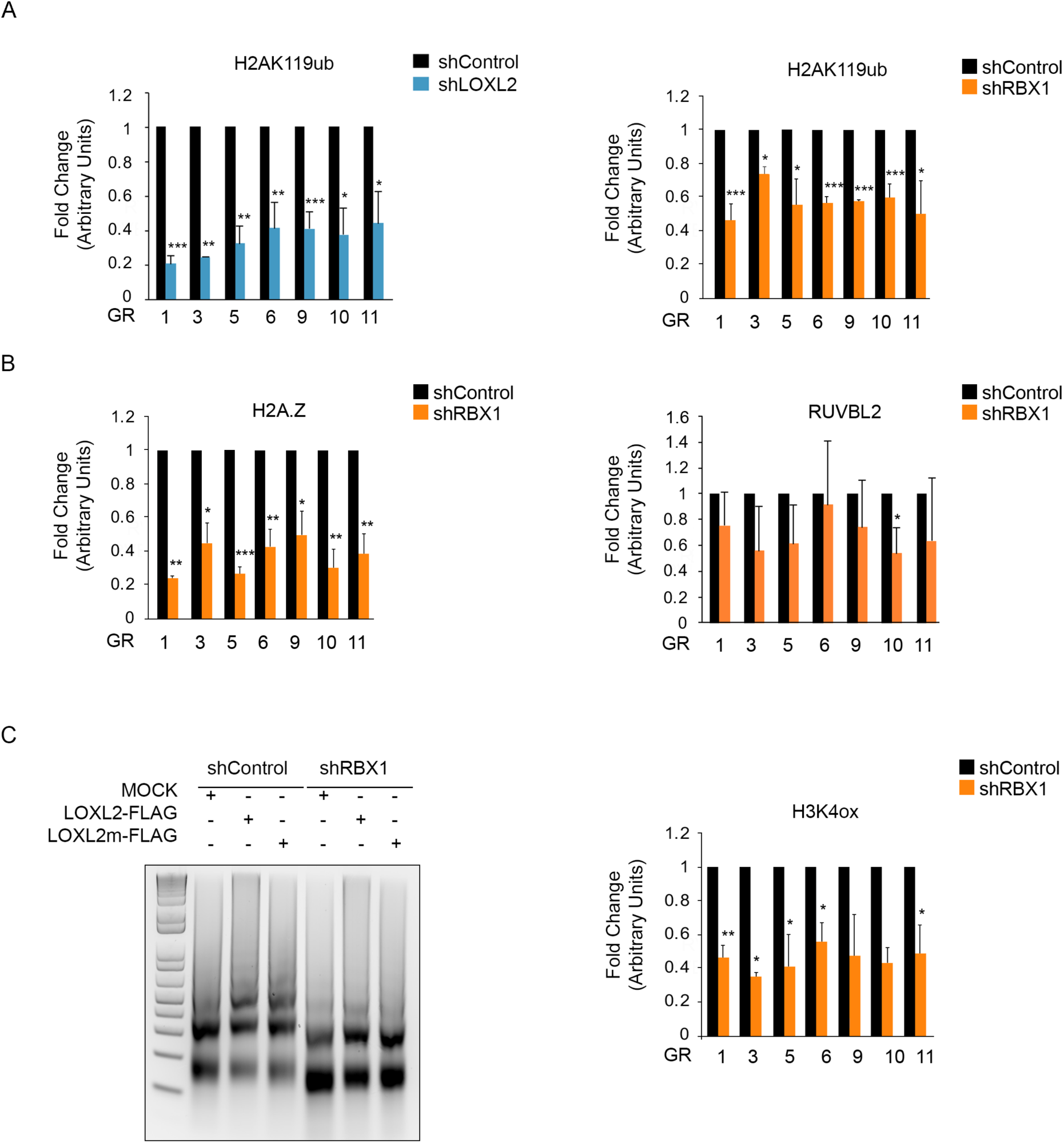
Ubiquitination of H2A is required for maintaining levels of H3K4ox and chromatin compaction. **A)** MDA-MB-231 cells were infected with shControl, shLOXL2, or shRBX1. ChIP-PCR experiments for H2AK119ub were performed in selected genomic regions (GR). Data of qPCR amplifications were normalized to the input and to total immunoprecipitated H2A. Results are expressed as a fold-change relative to data obtained shControl-cells, which were set to 1. Error bars indicate SD of at least three experiments * p < 0.05, ** p ≤ 0.01, *** p ≤ 0.001. **B)** MDA-MB-231 cells were infected with shControl or shRBX1. ChIP-PCR experiments of H2A.Z, RUVBL2 were performed in selected genomic regions (GR). Data of qPCR amplifications were normalized to the input and to total immunoprecipitated H3 for H3K4ox. Results are expressed as a fold change relative to the data obtained from shControl-cells, which were set to 1. **C)** HEK293T cells infected with shControl or shRBX1 were transfected with LOXL2 wild-type (LOXL2), inactive mutant LOXL2 (LOXL2m), or an empty vector (mock). At 48 hr post-transfection, isolated nuclei were digested with micrococcal nuclease (MNase) for 2 min, and total genomic DNA was analyzed using agarose gel electrophoresis (left panel). MDA-MB-231 cells were infected with shControl or shRBX1. ChIP-PCR experiments of H3K4ox were performed in selected genomic regions (GR). Data of qPCR amplifications were normalized to the input and to total immunoprecipitated H3 for H3K4ox. Results are expressed as a fold change relative to the data obtained from shControl-cells, which were set to 1 (right panel). Error bars indicate SD of at least three experiments * p < 0.05, ** p ≤ 0.01, *** p ≤ 0.001.

As heterochromatin properties depends on H3K9me3 levels (Peters et al., 2001), we analyzed H3K9me3 levels in all the conditions that caused a decrease in H3K4ox levels. ChIP-PCR in selected genomic regions and global levels of H3K9me3 were reduced in MDA-MB-231 cells knocked-down for LOXL2, RUVBL2 or RBX1 (Figure 7A). The same result was observed by ChIP-PCR for H3K9me2, but the levels of H3K27me3 remained constant (Figure S6). Finally, to assess globally the accesibility of chromatin, we performed ATAC-seq in the absence of RUVBL2, which previously was shown to cause a decrease in H3K4ox and H2A.Z levels. In cells with reduced levels of RUVBL2, no changes were observed in chromatin regions enriched in canonical euchromatin histone marks, such as H3K4me3 (Figure 7B, upper panel). Remarkably, more reads (indicative of open/accessible chromatin) were detected in those genomic regions decorated with heterochromatin histone marks of H3K9me3 and H3K27me3 (Figure 7B, lower panel). Interestingly, a similar pattern was observed in H3K4ox, H2A.Z and in common (H3K4ox, RUVBL2 and H2A.Z) regions (represented by 10213 peaks) in the absence of RUVBL2. Interestingly, this effect was not observed for RUVBL2-enriched regions.

**Figure 7.**
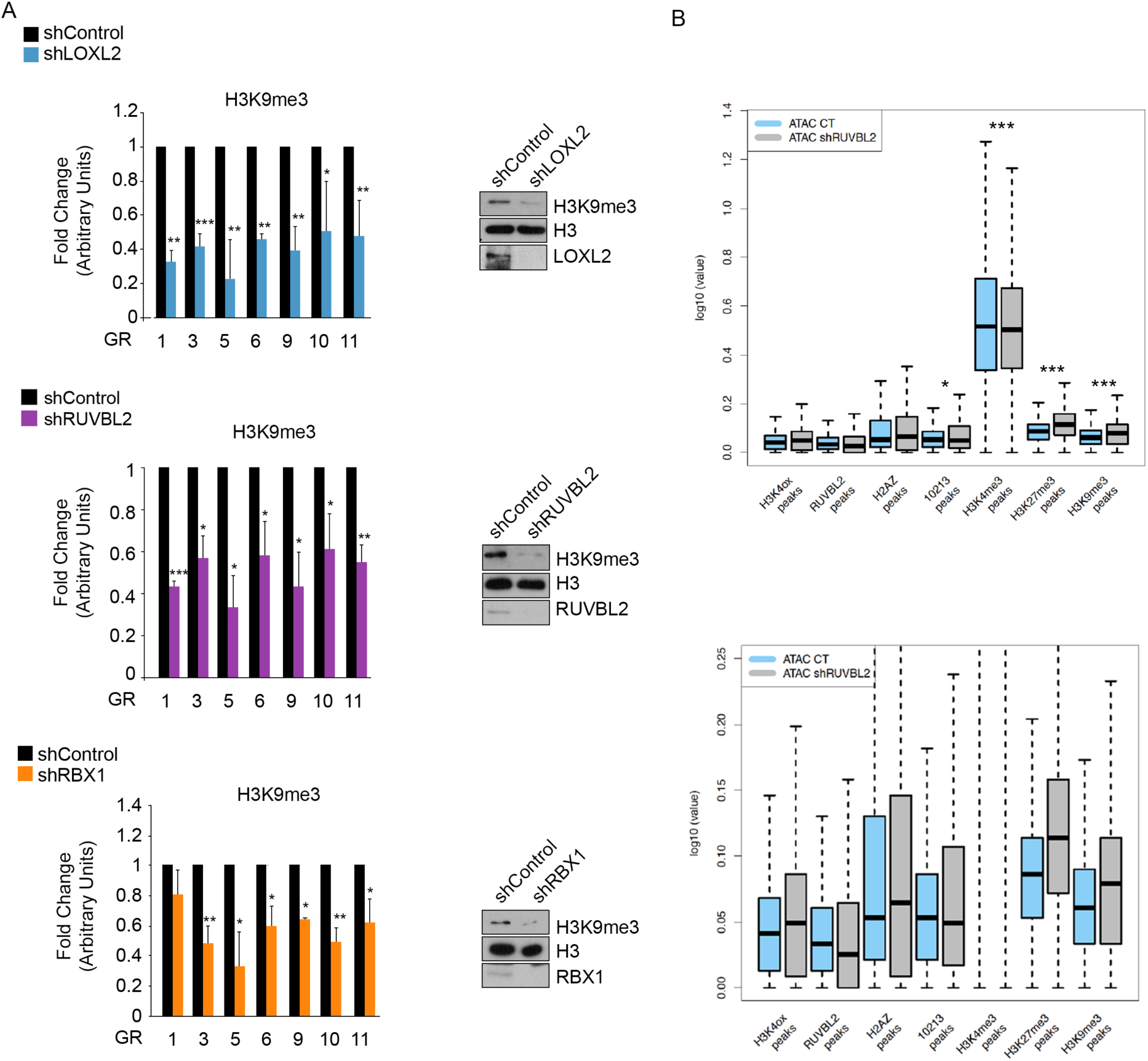
Maintaining H3K4ox levels is essential for maintaining H3K9me3 levels and chromatin compaction. **A)** MDA-MB-231 cells were infected with shControl, shLOXL2, shRUVBL2, or shRBX1. ChIP-PCR experiments of H3K9me3 were performed in the selected genomic regions (GR). Data of qPCR amplifications were normalized to the input and to total immunoprecipitated H3. Results are expressed as a fold-change relative to the data obtained from the shControl. Error bars indicate SD from at least three experiments *p < 0.05, **p ≤ 0.01, ***p ≤ 0.001. Total extracts were analyzed by Western blot using the indicated antibodies. **B)** ATAC-sequencing signals from different sets of ChIP-seq peaks in MDA-MB-231 cells infected with shControl or shRUVBL2. From left to right: i) H3K4ox, RUVBL2, H2A.Z, 10,213 peaks (common genomic regions with H3K4ox, RUVBL2 and H2A.Z), ii) H3K4me3 (ChIP from GSE107176), iii) H3K27me3 (ChIP from GSE107176), and iv) H3K9me3 (ChIP from GSE85158). The lower panel is a zoom of the upper one. *p ≤ 0.05, ***p ≤ 0.001.

### Heterochromatin Alterations Block the Oncogenic Properties of MDA-MB-231 Breast Cancer Cells

Based on the crucial role of heterochromatin in cell fitness, we next investigated if heterochromatin alteration could affect the oncogenic properties of the MDA-MB-231 TNBC cell line. For this purpose, MDA-MB-231 cells were knocked-down for each of the crucial members involved in the maintenance of H3K4ox (LOXL2, RUVBL2, DDB1, RBX1, or H2A.Z) and analyzed its effect in cell proliferation (using MTT) and colony formation assays (Figure S7A). All knock-downs were associated with a decrease in the proliferation rates (Figures S7B–7F) and reduced colony forming capacity compared to control cells (Figure 8A). Finally, reduced migration and invasion capacities were also observed in the same conditions (Figure 8B). To analyze whether restoring a normal heterochromatin state would also restore these oncogenic capacities in the knock-down lines, Suv39H1 (Lehnertz et al., 2003; Loyola et al., 2009) was reintroduced by retroviral infection in shLOXL2, shRUVBL2, or shRBX1 knocked-down MDA-MB-231 cells. Indeed, for each, we observed increased H3K9me3 levels (Figure S8A) and restored migration capacity (Figure 8C).

**Figure 8.**
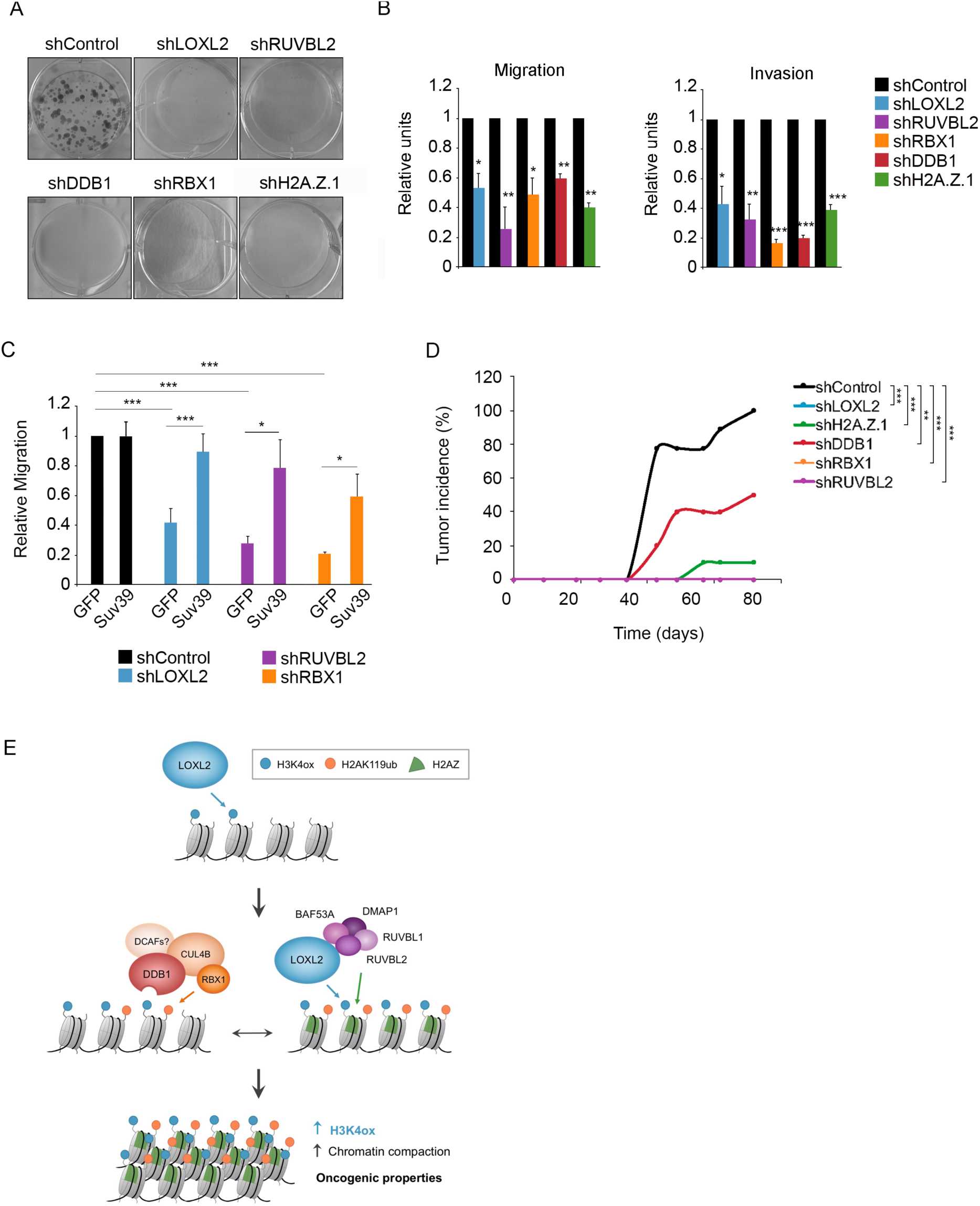
Opening heterochromatin blocks the oncogenic properties of MDA-MB-231 breast cancer cells. **A)** Colony-survival assay in MDA-MB-231 cells infected with shControl, shLOXL2, shRUVBL2, shDDB1, shRBX1, or shH2A.Z.1. **B)** Migration and invasion assays in MDA-MB-231 cells infected with shControl, shLOXL2, shRUVBL2, shDDB1, shRBX1, or shH2A.Z.1 after 96 hr of selection. Results are presented as the fold-change relative to the shControl cells, which were set to 1. **C)** MDA-MB-231 cells were first infected with shControl, shLOXL2, shRUVBL2, or shRBX1. After puromycin selection, knocked-down cells were re-infected with either GFP or SUV39H1-GFP and experiments were performed 48 hr later. Migration assays are presented as the fold-change relative to the shControl cells infected with GFP. In all panels, error bars indicate the SD from at least two independent experiments. *p < 0.05, **p ≤ 0.01, ***p ≤ 0.001. **D)** MDA-MB-231 cells were infected with shControl, shLOXL2, shDDB1, shH2A.Z.1, shRUVBL2, or shRBX1. After puromycin selection, 1×10^6^ knocked-down cells were injected into the mammary fat pad of nude mice. Tumor incidence was calculated as the percentage of tumors bigger than 100mm^3^ in different conditions and time points. Statistical analysis of day 72 comparing all cases with shControl is shown *p < 0.05, **p ≤ 0.01, ***p ≤ 0.001. **E)** Schematic representation of the proposed molecular mechanism.

To investigate the role of these components in tumor development *in vivo*, we implanted MDA-MB-231 cells infected with shLOXL2, shRUVBL2, shRBX1, shDDB1, shH2A.Z, or shControl, into the mammary fat pads of immunodeficient mice (Figure S8B) and tumor sizes were monitored for 10 weeks. Indeed, tumor growth was suppressed in shLOXL2, shRUVBL2, shRBX1, and shH2A.Z knock-downs. In shDDB1, reduced tumor incidence and a delay in tumor growth were observed (Figure 8D and S8C). Analysis of RNA levels of DDB1 in tumor samples at the end of the experiment showed that DDB1 knockdown was maintained (Figure S8D). Taken together, these data indicate that the maintenance of H3K4ox levels in chromatin promotes cell proliferation, invasion, and tumorigenesis both *in vitro* and *in vivo*.

Our finding showed that, in the TNBC MDA-MB-231 cell line, the epigenetic/nuclear function of LOXL2 in maintaining H3K4ox is critical for the cells’ tumorigenic/oncogenic capacities.

## DISCUSSION

Using an unbiased proteomic strategy, we found that LOXL2 interacts with the RUVBL1, RUVBL2, BAF53A, and DMAP1 proteins—all members of the SWI1-SNF2–related CBP activator protein (SRCAP) complex and the TIP60 chromatin remodeling complexes, both of which are involved in the deposition of the histone variant H2A.Z (Buschbeck and Hake, 2017; Huen et al., 2010; Morrison and Shen, 2009). H2A.Z.1 has been implicated in many cellular processes and plays a key role in heterochromatin establishment (Buschbeck and Hake, 2017; Rangasamy et al., 2003; Swaminathan et al., 2005).

We have demonstrated that RUVBL2 is essential for both maintaining the levels of H3K4ox and inducing chromatin compaction upon incorporation of H2A.Z into chromatin. Hence, both LOXL2 and RUVBL2 cooperate synergically to stably maintain their function as a writer and chromatin remodeler, respectively, which set the stage for consequent compaction of the chromatin structure (Figure 8F). The histone variant H2A.Z characteristically contains an extended acidic patch that leads to an altered nucleosome surface, and several pieces of evidence link this to chromatin compaction (Fan et al., 2002; Fan et al., 2004; Ryan and Tremethick, 2018; Zhou et al., 2007). Thus, we propose that incorporation of H2A.Z is essential for mediating changes in chromatin structure, favoring chromatin compaction, and maintaining H3K4ox levels. In fact, a significant increase of ATAC signal was observed in common H3K4ox, H2A.Z and RUVBL2-enriched regions. Moreover, in the absence of RUVBL2, the effect on chromatin compaction can be clearly observed in a subset of heterochromatin regions enriched with H3K9me3.

We successfully identified potential H3K4ox readers in nuclear extracts of the TNBC cell line MDA-MB-231, using a synthetically modified and biotinylated H3 peptide that resembles the intermediate product of the LOXL2 reaction. Although this is a fast reaction, we know from previous data that the intermediate alcohol is relatively stable and is maintained at least for two hours (Cebria-Costa et al., 2019). However, we cannot rule out that other proteins could also be readers of the lysine when it is completely oxidized. In fact, our in vivo experiments showed only a delay in tumor growth was observed when DDB1 was knocked-down, suggesting that other readers may exist. Using this approach, we identified that DDB1 is a reader of H3K4ox, and that it recruits the CRL4B complex that regulates the ubiquination of H2A. In line with this, several studies have demonstrated a crucial role of the CRL4B complex in creating repressive chromatin environments (Ji et al., 2014; Nakagawa and Xiong, 2011). Importantly, CRL4B is physically associated with SUV39H1, HP1, and DNMT3A, which facilitates H3K9me3 and DNA methylation, and is important for maintaining these heterochromatin features (Yang et al., 2015). In agreement with these functions, an orthologous complex of the human CRL4 in yeast, known as Cul4-Rik1, is required for heterochromatin formation and H3K9me3 (Jia et al., 2005; Kuscu et al., 2014). This complex contains the RING-box protein Pip1, cullin Cul4 (or Pcu4), and Rik1, which shares 21% of identity with DDB1 (Jackson and Xiong, 2009). Finally, our results strongly demonstrate that, acting through H2AK119ub, CRL4B is required for maintaining H2A.Z and H3K4ox levels in heterochromatin. This fits with previous evidence that post-translational modifications in histone tails can affect the incorporation of the histone variant H2A.Z (Choi et al., 2009; Raisner et al., 2005)

We also have shown that the described molecular mechanism is crucial for maintaining heterochromatin integrity. In particular, alteration of LOXL2, RUVBL2, or RBX1 dramatically affects global levels of H3K9me3 as well as the maintenance of H3K9me2/3 in heterochromatin regions. A slightly decrease in the RNA levels for SUV39H1/2 were observed in the RNAseq experiments when LOXL2 and RUVBL2 were knocked-down. Notably, the SUV39H1 methyltransferase is known to interact with the CRL4B complex (Yang et al., 2015) as well as with Snail1, a partner of LOXL2 that could bring it to heterochromatin (Dong et al., 2013; Millanes-Romero et al., 2013). These mechanisms could explain the decrease in H3K9me3 levels, when H3K4ox is reduced.

Remarkably, we demonstrated that the maintenance of heterochromatin domains through the LOXL2 function is essential for the oncogenic properties of TNBC cells. Consistent with these results, LOXL2 has been found to be overexpressed in many cancers and to be positively correlated with the acquisition of cellular malignancy and tumor formation (Fong et al., 2007; Martin et al., 2015; Moreno-Bueno et al., 2011; Peinado et al., 2008; Torres et al., 2015; Wong et al., 2014). In breast cancer, LOXL2 expression levels are higher in invasive and metastatic cell lines and patient-derived xenografts (PDXs) from triple negative breast patients (Cebria-Costa et al., 2019; Hollosi et al., 2009; Kirschmann et al., 2002). The *in vivo* evidence that shows how LOXL2 contribute to tumor progression and metastasis has been recently published (Salvador et al., 2017). Moreover, both its expression and its perinuclear localization is associated with a basal-like phenotype (Moreno-Bueno et al., 2011). Importantly, heterochromatin is mainly located at the nuclear periphery (Fawcett, 1966; Fedorova and Zink, 2008; Poleshko and Katz, 2014), which is consistent with the perinuclear localization of LOXL2 published before (Moreno-Bueno et al., 2011) and the enrichment of H3K4ox in heterochromatin. Both high LOXL2 levels and its perinuclear localization are associated with poor outcome and lower survival rates of patients with breast cancer (Ahn et al., 2013; Moreno-Bueno et al., 2011). Furthermore, both RUVBL2 and CRL4B complex are also known to play important roles in cancer biology (Grigoletto et al., 2011; Grigoletto et al., 2013; Jang et al., 2018; Mao and Houry, 2017).

The mechanism we describe here for maintaining H3K4ox in chromatin as an essential prerequisite for the oncogenic properties of breast cancer cells does not directly implicate regulation of gene expression. Specifically, dysregulated genes in shLOXL2- and shRUVBL2-knockdown conditions were not physically located in chromatin regions that normally contain H3K4ox and RUVBL2, and further, we can partially rescue the invasive and migrative phenotype by overexpressing Suv39H1. Therefore, we propose that these H3K4ox-enriched chromatin domains play more of a structural role that is important for establishing a proper balance between heterochromatin and euchromatin. This chromatin structure could be crucial for conferring other advantages to cancer cells independent of a direct effect on gene transcription regulation. For instance (and as we have previously suggested; (Cebria-Costa et al., 2019), it could protect cells from aberrant activation of the DNA damage response and any consequent cell cycle arrest or cell fate decisions, such as apoptosis or senescence. In fact, another LOX family member, LOXL3, has a major role in the maintenance of genomic stability (Santamaria et al., 2018). We also have demonstrated here that maintenance of this heterochromatin status is required for the ability of TNBC cells to migrate and invade. In agreement with this, several studies have shown that chromatin compaction plays a role in the migration process itself, regulating the mechanical properties of the nucleus (Fu et al., 2012; Gerlitz and Bustin, 2010; Gerlitz and Bustin, 2011; Gerlitz et al., 2007; Yokoyama et al., 2013). Accordingly, migration ability is strongly impaired after overexpressing a dominant-negative form of histone H1 or of SUV39H1, as well as after treatment with chemical compounds that promote an open chromatin state (Fu et al., 2012; Gerlitz and Bustin, 2010; Gerlitz et al., 2007).

Our experiments revealed a direct link between the maintenance of H3K4ox levels, chromatin condensation, and tumorigenic capacities of a cancer cell. Thus, these data provide novel therapeutic opportunities and suggest that any of the components described could be a targetable key for establishing effective cancer therapy in TNBC tumors, which are often associated with therapy resistance and poor prognosis. Along that line, one actionable target could be the link between chromatin compaction and cancer progression: using drugs that reduce chromatin compaction and generate an open chromatin state could increase DDR signaling and severely reduce both the oncogenic potential and survival rates of cancer cells (Cebria-Costa et al., 2019; Di Micco et al., 2011). Knowing the mechanistic details behind the role of H3K4ox will now set the stage for developing such new strategies.

## ACKNOWLEDGMENTS

We would like to thank R. Peña for technical assistance, V.A. Raker for manuscript editing, T. Jenuwein for SUV-39 constructs, Jean Rosenbaum for RUVBL2 constructs, and Carla Nieto and Rita Webster for their experimental support. Proteomics were performed in the CRG/UPF Proteomics Unit, which is part of the of Proteored, PRB3, and is supported by grant PT17/0019, of the PE I+D+i 2013-2016, funded by ISCIII and ERDF. SNAP experiments were done at the IRB in The Advanced Digital Microscopy (ADM) Core Facility. This work was supported by grants from Instituto de Salud Carlos III (ISCIII) FIS/FEDER (PI12/01250; CP08/00223; PI16/00253 and CB16/12/00449), MINECO, FPU14/04071 to G.S and Juan de la Cierva Incorporation fellowship IJCI-2014-20723 to G.V. MICIU (PGC2018-095616-B-I00/GINDATA and FEDER) to T.H.S., Breast Cancer Research Foundation (BCRF-17-008) to J.A., and Red Temática de Investigación Cooperativa en Cáncer (RTICC, RD012/0036/005) to J.A. T.H.S. was supported by institutional funding (MINECO) through the Centres of Excellence Severo Ochoa award and the CERCA Program of the Catalan Government, and S.S.B. was supported by a Fundació La Caixa fellowship and C.J. was supported by EMBO and CIHR fellowships. We thank the FERO Foundation, La Caixa Foundation for financial support (LCF/PR/PR12/51070001) and Cellex Foundation for providing research facilities and equipment.

## Author Contributions

S.P. developed the study concept; G.S. and S.P. directed experimental design and interpreted the data; E.B, G.V. and R.A.C. performed computational analysis; T.T., L.P., J.Q., A.M., and C.J. conducted additional experiments; S.S. and T.H.S. helped with SNAP experiments; P.N. and S.S. were the pathologists involved in LOXL2 immunohistochemistry analysis; C.V. and R.D. conducted analyses; V.P. and C.S. performed sample and data acquisition; B.M, ME and J.A. carried out *in vivo* experiments; J.V. performed the proteomic analyses; and L.D.C. and A.G.H. interpreted data and helped formulate the discussion. All authors read, edited, and approved the manuscript.

## Declaration of Interests

The authors declare no competing interests.

## Data and materials availability

GSE142463

## MATERIALS AND METHODS

### Cell Lines and Culture Conditions

HEK293T cells (ATCC No.: CRL-3216) and MDA-MB-231 cells (ATCC No.: HTB-26) were cultured in Dulbecco’s modified Eagle’s medium (Biowest; L0106-500) supplemented with 1% penicillin/streptomycin (Gibco; 15140122), 2 mM L-glutamine (Biowest; X0550-100), and 10% FBS (Gibco; 10270106) at 37°C in 5% CO_2_. The lack of mycoplasma contamination was tested regularly using standard PCR with the primers:

F: 5′-GGCGAATGGGTGAGTAACACG-3′

R: 5′-CGGATAACGCTTGCGACCTATG-3′

### Transfection

For overexpression assays, HEK293T cells were seeded for 18–24 hr and transiently transfected with the indicated vectors using either the JetPrime reagent (Polyplus-transfection; 114-15) or polyethylenimine polymer (Polysciences Inc; 23966-1), following manufacturer’s instructions.

### Lentivirus Production and Infection

For lentiviral infections, HEK293T cells were used to produce viral particles. Cells were grown to 70% confluence (day 0) and transfected by adding (dropwise) a mixture of 150 mM NaCl, DNA (50% of the indicated shRNA vector, 10% pCMV-VSVG, 30% pMDLg/pRRE, and 10% pRSV rev), and polyethylenimine polymer (Polysciences Inc; 23966-1), which had been pre-incubated for 15 min at room temperature. The transfection medium was replaced with fresh medium after 24 hr (day 1). On days 2 and 3, the cell-conditioned medium was filtered with a 0.45 µm filter unit (Merk Millipore; 051338) and stored at 4°C. Viral particles were concentrated with a Lenti-X Concentrator (Clonetech; 631232) following manufacturer’s instructions, and virus aliquots were stored at –80°C until use. HEK293T and MDA-MB-231 cells were infected by incubating with concentrated viral particles for 16–18 hr, and then medium was changed for fresh medium containing 2.7 µg/ml or 2.5 µg/ml puromycin, respectively. After 48 hr of selection, cells were seeded, and experiments were performed at 48 hr post-selection.

### Retrovirus Production and Infection

For retroviral infections, HEK293T cells were transfected with pCL-ampho packaging vector and the indicated vectors using JetPrime reagent (Polyplus-transfection; 114-15) following manufacturer’s instructions (day 0). The transfection medium was replaced with fresh medium after 24 hr, and cell-conditioned medium at days 2 and 3 was filtered and used to infect MDA-MB-231 cells twice. Target cells were seeded into a 6-well plate and infected by adding the filtered retroviral supernatant and centrifuging the plates at 32°C for 2 hr at 1,000 × g. After that, 1 ml of fresh complete medium was added, and cells were incubated at 37°C.

### Ubiquitination Assays

To study ubiquitination, the DUB inhibitor PR-619 (Tocris Bioscience; 4482) was added to lysis buffers to 100 µM, to help stabilize the ubiquitinated proteins.

### Total Extracts

Cells were lysed in SDS lysis buffer (2% SDS, 50 mM Tris-HCl pH [7.5], 10% glycerol). Samples were kept at room temperature to avoid SDS precipitation and passed through a syringe to homogenize them. Proteins were quantified with the DC Protein Array kit (Lowry method; Bio-Rad) and Nanodrop analysis.

### Histone isolation

Cells were pelleted by centrifugation and washed with cold PBS. Pellets were resuspended by vortexing with lysis buffer (10 mM Tris, pH 6.5, 50 mM sodium bisulfite, 1% Triton X-100, 10 mM MgCl_2_, 8.6% sucrose, 10 mM sodium butyrate) and centrifuged at full speed for 15 seconds twice. The same procedure was repeated once with wash buffer (10 mM Tris [pH 7.4], 13 mM EDTA). Pellets were resuspended in 0.4 N H_2_S0_4_ and left for 1 hr at 4 °C with occasional gentle mixing (by hand). After centrifuging at full speed for 5 min, supernatants were transferred to a new tube, and acetone was added (1:9). The mixture was left overnight at –20 °C and then centrifuged at full speed for 10 min. Pellets were air dried for 5 min and resuspended in 30–100 µl of sterile water. The protein level was quantified with the DC Protein Array kit (Lowry method; Bio-Rad) before sample preparation.

### Western Blots

Western blots were performed according to standard procedures. Briefly, samples were mixed with 5× loading buffer (250 mM Tris-HCl [pH 6.8], 10% SDS, 0.02% Bromophenol blue, 50% glycerol, 20% β-mercaptoethanol) and boiled at 95°C for 5 min. Proteins were analyzed by sodium dodecyl sulfate polyacrylamide gel electrophoresis (SDS-PAGE) using different percentages of polyacrylamide concentrations, ranging from 7.5% to 15%. Gels were run in Tris-glycine-SDS (TGS) buffer (25 mM Tris-OH [pH 8.3], 192 mM glycine, and 5% SDS), and proteins were transferred to a nitrocellulose membrane (Amersham Protran 0.45 nitrocellulose, GE Healthcare) in transfer buffer (50 mM Tris-OH, 396 mM glycine, 0.1% SDS, and 20% methanol) for 60 to 120 min, depending on the molecular weight of the protein. Membranes were stained with Ponceau S (0.5% Ponceau S and 1% acetic acid) and blocked with 5% non-fat milk or bovine serum albumin (BSA) in TBS-T buffer (25 mM Tris-HCl [pH 7.5], 137 mM NaCl, and 0.1% Tween) for 1 hr. Primary antibodies were added in fresh blocking solution and incubated overnight at 4 °C. After three washes of 10 min with TBS-T, membranes were incubated with horseradish peroxidase (HRP)-combined secondary antibodies in fresh blocking solution for 1 hr at room temperature. After a new round of washes, membranes were developed by incubation with a substrate for HRP-enhanced chemiluminiscence (ECL) and exposure to autoradiography films. To detect low-quantity (or otherwise difficult-to-detect) proteins, more sensitive ECL and films were used.

### Co-immunoprecipitation (co-IP) Assays

HEK293T cells were transfected with pcDNA3-hLOXL2wt-Flag or an empty pcDNA3. After 48 hr, co-immunoprecipitation assays were carried out as previously described (Herranz et al., 2016) with the following modifications. Cells were washed with 1× PBS pre-warmed to 37°C and incubated first with 1 mM DTBP solution for 30 min at 37 °C and then with cold 100 mM Tris-HCl (pH 7.4) for 5 min on ice. Cells were washed once with cold PBS, lysed in high-salt lysis buffer (20 mM HEPES [pH 7.4], 10% glycerol, 350 mM NaCl, 1 mM MgCl_2_, 0.5% Triton X-100, and 1 mM dithiothreitol) with protease inhibitors, and incubated for 30 min on ice. Lysate samples were centrifuged at 13,000 rpm at 4°C for 10 min. Balance buffer (20 mM HEPES [pH 7.4], 1 mM MgCl_2_, and 10 mM KCl) was added to the resulting supernatant to reach a final NaCl concentration of 150 mM. Cell extracts were incubated with Flag-M2 Affinity Agarose Gel (Sigma-Aldrich; A2220) for 4 hr at 4°C and washed three times with wash buffer (20 mM HEPES [pH 7.4], 1 mM MgCl_2_, 150 mM NaCl, 10% glycerol, 0.5% Triton X-100). Precipitated complexes were eluted with 2× loading buffer.

For endogenous co-IP assays in MDA-MB-231 cells, DTBP treatment was performed as described before, and cells were lysed with soft lysis buffer (50 mM Tris [pH 8], 10 mM EDTA, 0.1% NP-40, and 10% glycerol) supplemented with protease and phosphatase inhibitors. After incubation for 5 min on ice, lysates were centrifuged at 3,000 rpm at 4°C for 15 min, and pellets were lysed with high-salt lysis buffer and balanced as explained above. Antibodies were added for 16–18 hr at 4°C, followed by immunoprecipation of immunocomplexes with protein-A agarose beads (Diagenode; C03020002) previously blocked with 0.5% BSA for 1 hr at 4°C. Immunoprecipitated complexes were washed three times with wash buffer and eluted with 2× loading buffer. When used, ethidium bromide was added in lysis and washing buffers (100 µg/ml).

### Nucleosome purification and oxidation assays

Nucleosomes were purified from HEK293T cells. After harvesting by centrifugation, cells were resuspended in buffer A (10 mM Tris buffer [pH 7.4], 10 mM NaCl, 3 mM MgCl_2_, 300 mM sucrose, 0.2% NP-40) supplemented with protease inhibitors and homogenized by ten strokes in a Dounce homogenizer. Cells were then centrifuged, and nuclei pellets were resuspended in buffer A with 10 mM CaCl_2._ Four units of micrococcal nuclease were added for 30 min at room temperature, and the reaction was stopped by the addition of 50 mM EDTA. Purified nucleosomes were aliquoted and kept at –80°C. HEK293T cells were transfected with pcDNA3-hLOXL2wt-Flag or an empty pcDNA3. After 48 hr, co-immunoprecipitation assays were performed as described before but without DTBP treatment. After incubation with Flag-M2 Affinity Agarose Gel (Sigma-Aldrich; A2220), precipitated complexes were resuspended in oxidation buffer (20 mM HEPES [pH 7.4], 100 mM NaCl, 1 mM MgCl_2_, 0.1 mg/ml BSA, 1 mM DTT) and incubated with 50 ng of purified nucleosomes for 2 hr at 37°C. Then, 5× loading buffer was added to samples, and proteins were analyzed by SDS-PAGE and Western blotting.

### Readers experiment

MDA-MB-231 cells were lysed in soft lysis buffer (50 mM Tris [pH 8], 10 mM EDTA, 0.1% NP-40, 10% glycerol) supplemented with protease inhibitors for 5 min on ice and centrifuged at 3,000 rpm for 15 min. The supernatant was discarded, and the nuclear pellet was resuspended in high-salt lysis buffer (20 mM HEPES [pH 7.4], 10% glycerol, 350 mM NaCl, 1 mM MgCl_2_, 0.5% Triton X-100, 1 mM dithiothreitol) for 30 min at 4°C. Samples were centrifuged at 13,000 rpm for 10 min and the NaCl concentration of the supernatant was reduced to 300 mM NaCl with balance buffer (20 mM HEPES [pH 7.4], 1 mM MgCl_2_, 10 mM KCl). To preclear the nuclear extracts, samples were incubated with streptavidin beads for 1-2 hr. Beads were separated, and nuclear extracts were incubated for 1.5 hr at 4°C with 1 µg of H3K4ox (ART-K(OH)-QTARKSTGGKAP-biotin) or irrelevant (AHIVMVDAYKPTK-NH(CH_2_)_2_NH-biotin) peptides that had been previously recovered with streptavidin magnetic beads (Invitrogen; 65305) for 30 min at 4°C. Finally, immunoprecipitated proteins were washed two times with wash buffer (20 mM HEPES [pH 7.4], 1 mM MgCl_2_, 150 mM NaCl, 10% glycerol, 0.5% Triton X-100) and once with high salt lysis buffer, and processed for mass spectrometry analysis.

### Sample preparation for mass spectrometry

Beads used in immunoprecipitation were cleaned three times with 500 µl of 200 mM ammonium bicarbonate and 60 µl of 6 M urea / 200 mM ammonium bicarbonate were added. Samples were then reduced with dithiothreitol (30 nmol, 37 °C, 60 min), alkylated in the dark with iodoacetamide (60 nmol, 25 °C, 30 min) and diluted to 1 M urea with 200 mM ammonium bicarbonate for trypsin digestion (1 µg, 37°C, 8 hr, Promega cat # V5113). After digestion, the peptide mix was acidified with formic acid and desalted with a MicroSpin C18 column (The Nest Group, Inc) prior to LC-MS/MS analysis.

### Chromatographic and mass spectrometric analysis

Samples were analyzed using a LTQ-Orbitrap Velos Pro mass spectrometer (Thermo Fisher Scientific, San Jose, CA, USA) coupled to an EASY-nLC 1000 (Thermo Fisher Scientific (Proxeon), Odense, Denmark). Peptides were loaded onto the 2-cm Nano Trap column with an inner diameter of 100 µm packed with C18 particles of 5 µm particle size (Thermo Fisher Scientific) and were separated by reversed-phase chromatography using a 25-cm column with an inner diameter of 75 µm, packed with 1.9 µm C18 particles (Nikkyo Technos Co., Ltd. Japan). Chromatographic gradients started at 93% buffer A and 7% buffer B with a flow rate of 250 nl/min for 5 min and gradually increased 65% buffer A and 35% buffer B in 60 min. After each analysis, the column was washed for 15 min with 10% buffer A and 90% buffer B. Buffer A: 0.1% formic acid in water. Buffer B: 0.1% formic acid in acetonitrile. The mass spectrometer was operated in positive ionization mode with nanospray voltage set at 2.1 kV and source temperature at 300°C. Ultramark 1621 was used for external calibration of the FT mass analyzer prior the analyses and an internal calibration was performed using the background polysiloxane ion signal at m/z 445.1200. The acquisition was performed in data-dependent adquisition (DDA) mode and full MS scans with 1 micro scans at resolution of 60,000 were used over a mass range of m/z 350-2000 with detection in the Orbitrap. Auto gain control (AGC) was set to 1E6, dynamic exclusion (60 seconds) and charge state filtering disqualifying singly charged peptides was activated. In each cycle of DDA analysis, following each survey scan, the top twenty most intense ions with multiple charged ions above a threshold ion count of 5000 were selected for fragmentation. Fragment ion spectra were produced via collision-induced dissociation (CID) at normalized collision energy of 35% and they were acquired in the ion trap mass analyzer. AGC was set to 1E4, isolation window of 2.0 m/z, an activation time of 10 ms and a maximum injection time of 100 ms were used. All data were acquired with Xcalibur software v2.2. Digested bovine serum albumin (New england biolabs cat # P8108S) was analyzed between each sample to avoid sample carryover and to assure stability of the instrument; QCloud was used to control instrument longitudinal performance during the project(Chiva et al., 2018).

### Proteomic Data Analysis

Acquired spectra were analyzed using the Proteome Discoverer software suite (v1.4, Thermo Fisher Scientific) and the Mascot search engine (v2.5 Matrix Science)(Perkins et al., 1999). The data was searched against a Swiss-Prot human database (as in April 2017, 20797 entries) plus a list of common contaminants and all the corresponding decoy entries(Beer et al., 2017). For peptide identification, a precursor ion mass tolerance of 7 ppm was used for MS1 level, trypsin was chosen as enzyme, and up to three missed cleavages were allowed. The fragment ion mass tolerance was set to 0.5 Da for MS2 spectra. Oxidation of methionine and N-terminal protein acetylation were used as variable modifications whereas carbamidomethylation on cysteines was set as a fixed modification. False discovery rate (FDR) in peptide identification was set to a maximum of 5%.

### Interactome statistical analysis

The inferential statistical analysis was done using the open-source statistical package R. Files containing all spectral counts for each sample and its replicates were imported into the R software from the result tables of Proteome Discoverer (v1.4, Thermo Fisher Scientific). Data were assembled into a matrix of spectral counts, in which columns represented the different conditions and rows represented the identified proteins (Gregori et al., 2012). An unsupervised exploratory data analysis (EDA) consisting of data normalizations, principal components analysis, and hierarchical clustering of the samples on the SpC matrix was first performed. The Generalized Linear Model based on the Poisson distribution was used as a significance test (Gregori et al., 2013). Finally, the Benjamini and Hochberg multitest correction was used to adjust the p-values to the control of the FDR. To identify statistically significant proteins, spectral count signal, fold change, and adjusted p-value were taken into account (Gregori et al., 2013).

### Dot blot assay

To check that the peptides were equally recovered, in parallel with the experiment, an additional microgram of H3K4ox or irrelevant peptides were incubated with streptavidin beads and eluted with 20 µl of SDS 1%. Then, 5× loading buffer was added and samples were boiled at 95°C for 5 min. For dot blot assays, 2–10 µl of each peptide were applied to a nitrocellulose membrane freehand. The blot was blocked in 5% milk in TBS-T for 30 min to 1 hr at room temperature and incubated with anti-biotin overnight at 4°C. Secondary antibody incubation and developing was performed as described for Western blots.

### Pulldown Assays

H3K4ox (ART-K(OH)-QTARKSTGGKAP-biotin) or irrelevant (AHIVMVDAYKPTK-NH(CH_2_)_2_NH-biotin) peptides (1 µg) were immobilized onto magnetic streptavidin beads (previously blocked with 100 µg/ml BSA) in binding buffer (50 mM Tris-HCl [pH 8], 150 mM NaCl and 100 µg/ml BSA) for 30 min at 4 °C. Samples were then washed with binding buffer and incubated with 2 µg of DDB1-GST or GST for 1 hr at 4°C. Complexes were washed successively once with wash buffer 1 (50 mM Tris-HCl [pH 8], 150 mM NaCl, 0.1% Triton), twice with wash buffer 2 (50 mM Tris-HCl [pH 8], 300 mM NaCl, 0.1% Triton), and a final time with wash buffer 1. Proteins bound to streptavidin beads were resuspended in loading buffer and analyzed by Western blot. Recovery of peptides was analyzed by Tricine SDS-PAGE as previously described (Haider et al., 2012), transferred to a nitrocellulose membrane, and incubated with streptavinidin-HRP.

### Chromatin association assay

Cells were crosslinked with 1% formaldehyde in PBS for 10 min at 24°C, and pellets were resuspended in buffer A (100 mM Tris, pH [7.5], 5 mM MgCl_2_, 60 mM KCl, 125 mM NaCl, 300 mM sucrose, 1% NP-40, 0.5 mM DTT). After incubating 10 min on ice, samples were centrifuged and lysed in buffer B (3 mM EDTA, 0.2 mM EGTA, 1 mM DTT) twice. The chromatin-containing pellets were resuspended in buffer C (1% SDS, 10 mM EDTA, 50 mM Tris [pH 8.0]) overnight at 4°C. Samples were centrifuged 2 min at 16,100 × g, and the supernatant was quantified and used for Western blotting.

### Subcellular fractionation assay

Subcellular fractionation assays were performed as previously described (Wang et al., 2006). Cells were harvested in hypotonic buffer (10 mM Tris-HCl [pH 7.4], 10 mM KCl, 1.5 mM MgCl_2_, 0.2 mM PMSF, 1 mM DTT, and protease inhibitors), incubated for 20 min on ice and disrupted by homogenization in a douncer 10 times. Samples were centrifuged for 10 min at 3,600 rpm, and the supernatant was saved as the cytoplasmic fraction. Pellets were resuspended in extraction buffer (15 mM Tris-HCl [pH 7.4], 0.4M NaCl, 1 mM MgCl2, 1 mM EDTA, 10% glycerol, 0.2 mM PMSF, 1 mM DTT, and protease inhibitors), incubated for 30 min by rotating at 4 °C, and centrifuged at 13,200 rpm for 30 min. The supernatant was saved as the nuclear extract and the pellet, which was the chromatin fraction, was resuspended in SDS lysis buffer (2% SDS, 50 mM Tris-HCl [pH 7.5], 10% glycerol).

### Chromatin Immunoprecipitation (ChIP)

Cells were crosslinked in 1% formaldehyde for 10 min at 37⁰C and then glycine was added to a final concentration of 0.125 M for 2 min at room temperature to quench the crosslinking. Cells were scraped with cold soft-lysis buffer (50 mM Tris-HCl [pH 8], 10 mM EDTA, 0.1% NP-40, and 10% glycerol) supplemented with protease inhibitors, and samples were centrifuged at 3,000 rpm for 15 min. Nuclei pellets were lysed with SDS-lysis buffer (50 mM Tris-HCl [pH 8], 1% SDS, and 10 mM EDTA) supplemented with protease inhibitors and sonicated to generate 200–600 bp DNA fragments. After 20 min of incubation on ice, sonicated extracts were centrifuged 10 min at 13,000 rpm and supernatants were diluted 1:10 with dilution buffer (0.01% SDS, 1.1% Triton X-100, 1.2 mM EDTA, 16.7 mM Tris-HCl [pH 8], and 167 mM NaCl).

For immunoprecipitation, samples were incubated by rotating overnight at 4°C with primary antibody or irrelevant IgGs and then incubated with protein-A agarose beads (Diagenode; C03020002) previously blocked with 0.5% BSA for 3 hr at 4 °C with rotation. The bead slurry was sequentially washed three times with low-salt buffer (0.1% SDS, 1% Triton X-100, 2 mM EDTA, 20 mM Tris-HCl [pH 8], and 150 mM NaCl), three times with high-salt buffer (as the low-salt but with 500 mM NaCl), and twice with LiCl buffer (250 mM LiCl, 1% Nonidet P-40, 1% sodium deoxycholate, 1 mM EDTA, and 10 mM Tris-HCl [pH 8]). Beads were then incubated with elution buffer for 1 hr at 37 °C and overnight at 65°C with 200 mM NaCl to reverse formaldehyde crosslinking. Finally, samples were treated for 1 hr at 55⁰C with proteinase K solution (0.4 mg/mL proteinase K (Roche; 3115828001), 50 mM EDTA, and 200 mM Tris [pH 6.5]). DNA was purified with MinElute PCR purification kit (Qiagen; 28006) and eluted in MilliQ water (80–100 µl). The amount of immunoprecipitated DNA was analyzed by quantitative PCR (qPCR) in duplicates or triplicates; in a final volume of 10 µl, 4 µl of the eluted DNA was mixed with the forward and reverse primers (Sigma; 100-500 nM each) and 1× PerfeCTa® SYBR® Green FastMix (Quantabio; 95073-01). Thermocycling parameters used were: 95 °C 30 s; 40 cycles 95 °C 5 s, 60 °C 15 s, 72 °C 10s; melting curve. ChIP results were quantified relative to the input amount.

### ChIP-seq

For ChIP sequencing (ChIP-seq) analysis, the NEBNext Ultra DNA library Prep Kit for Illumina was used to prepare the libraries, and samples were sequenced using the Illumina HiSeq 2500 system.

### Analysis of ChIP-seq Data

ChIP-seq samples were mapped against the hg19 human genome assembly using Bowtie with default parameters. In particular, the option –m must be off to allow for the detection of multi-locus reads that are mapped into more than one region (Langmead et al., 2009). MACS was run with the default parameters but with the shift-size adjusted to 100 bp to perform the peak calling against the corresponding Input sample (Zhang et al., 2008). To build the final set of 10,213 H3K4ox regions furnished with other features, we firstly intersected the peaks reported in common between both replicates of H3K4ox. Next, H3K4ox peaks reported in both replicates were overlapped with the peaks previously reported for RUVBL2 and H2A.Z. The set of 2,267 genes was retrieved by matching the 10,213 peaks with the full collection of RefSeq gene transcripts (O’Leary et al., 2016). Reports of functional enrichments of GO and other genomic libraries were generated using the EnrichR tool (Kuleshov et al., 2016). The heatmaps displaying the density of ChIP-seq reads around the summit of each ChIP-seq peak were generated by counting the number of reads in this region +/- 5 kbp for each individual peak and normalizing this value with the total number of mapped reads of each sample. Peaks on each ChIP heatmap were ranked by the logarithm of the average number of reads in the same genomic region. The UCSC genome browser was used to generate the screenshots of each group of experiments (Kent et al., 2002).

### MNase Assay

HEK293T cells infected with shRNAs (as indicated) and selected for 48 hr with puromycin were seeded and transfected with pcDNA3-empty or pcDNA3-hLOXL2wt/mut-Flag. At 48-hr post-transfection, 1.5 × 10^6^ cells per condition were used for the MNase assay. Cell pellets were lysed in 500 µL of buffer A (10 mM Tris [pH 7.4], 10 mM NaCl, 3 mM MgCl_2_, and 0.3 M sacarose) supplemented with protease inhibitors for 10 min at 4⁰C. NP-40 was added to a final concentration of 0.2% (v/v) and incubated for 10 min at 4 °C. The lysate was centrifuged for 10 min at 1200 rpm at 4 °C, and the resulted pellet was resuspended in 100 µl of buffer A containing CaCl_2_ to a final concentration of 10 mM. MNase digestion was carried out with 0.004 units for 2 min at room temperature, and the enzyme was inactivated with 50 mM EDTA. Finally, extracts were treated with RNase A for 2 min at room temperature followed by proteinase K for 10 min at 56 °C. DNA was purified using MinElute PCR purification kit (Qiagen; 28006), and digestion products were analyzed by 1.5% agarose gel electrophoresis using Gelred TM gel stain (Biotium; BT-41003) for DNA visualization.

### ATAC Sequencing

MDA-MB-231 cells were infected with lentiviral particles for shControl and shRUVBL2, selected for 48h with puromycin and 48 hr post-selection ATAC-seq experiment was performed as described (Buenrostro et al., 2013). About 50,000 cells of each condition were treated with transposase Tn5 (Nextera DNA Library Preparation Kit, Illumina), and DNA was purified using MinElute PCR Purification Kit (Qiagen; 28006). Transposed DNA was amplified by PCR using NEBNextHigh-Fidelity 2× PCR Master Mix (New Englands Labs; M0541S) and primers containing a barcode. The number of cycles for library amplification was calculated as described (Buenrostro et al., 2013). DNA was again purified using MinElute PCR Purification kit, and samples were sequenced using the Illumina HiSeq 2500 system.

### Analysis of ATAC-seq Data

ATAC-seq samples were mapped against the hg19 human genome assembly using Bowtie with the option –m 1 to discard those reads that could not be uniquely mapped to just one region, and with the option –X 2000 to define the maximum insert size for paired-end alignment (Langmead et al., 2009). Mitochondrial reads were removed from each resulting map and down-sampling was used to obtain the same number of mapped fragments per sample. Boxplots showing the ATAC-seq level distribution for a particular experiment on a set of ChIP-seq genomic peaks were calculated by determining the maximum value on each peak at this sample normalized by the total number of reads.

### RNA extraction

Cells were washed three times with PBS and lysed with 800 µl of TRIzol® reagent (Invitrogen; 15596018). 200 µl of RNase-free chloroform were added, mixed and incubated at room temperature for 2 min. The solution was centrifuged at 12,000 × g for 15 min at 4°C and the upper aqueous phase was transferred to a new tube. 500 µl of RNase-free isopropanol were added and solution was incubated for 10 min at room temperature. RNA was then precipitated by centrifugation at 12,000 × g for 10 min at 4 °C. Pellets were washed once with 1 ml of 75% RNase-free ethanol and centrifuged at 7,500 × g for 5 min at 4 °C. RNA pellets were air-dried for 5–10 min to eliminate ethanol, resuspended in DEPC water, and dissolved for 10 min at 60 °C. RNA was quantified with Nanodrop.

### Quantitative RT-PCR

RNA was retrotranscribed using iScriptTM Reverse Transcription Supermix (Biorad; 1708841) following manufacturer’s instructions. Quantitative determination of RNA levels was performed in duplicate or triplicate in a final volume of 10 µl with 15–100 ng of cDNA, forward and reverse primers (Sigma; 100-500 nM each) and 1× PerfeCTa® SYBR® Green FastMix (Quantabio; 95073-01). Thermocycling parameters used were: 95 °C 30 s; 40 cycles 95 °C 5 s, 60 °C 15 s, 72°C 10 s; melting curve. Values were normalized to the expression of housekeeping genes (*HPRT* or *Pumilio*).

### RNA Sequencing

MDA-MB-231 cells were infected with lentiviral particles for shControl and shRUVBL2. At 48 hr post-selection, RNA was extracted with an RNeasy Mini Kit (Qiagen; 74104) following manufacturer’s instructions. RNA-seq experiments were performed with two biological replicates of each condition and samples were sequenced using the Illumina HiSeq 2500 system.

### Analysis of RNA-seq Data

RNA-seq samples were mapped against the hg19 human genome assembly using TopHat (Trapnell et al., 2009). Cuffinks was run to quantify the expression in FPKMs of each annotated transcript in RefSeq and to identify the list of differentially expressed genes for each case (FDR ≤ 0.05 and log_2_ FC ≥ 0.58) (Trapnell et al., 2012). Gene ontology analyses of the deregulated genes were generated using the EnrichR tool (Kuleshov et al., 2016).

### SNAP-based experiments

MDA-MB-231 cells stably expressing SNAP-tag histones were generated by infection with retroviral particles containing pBabe-H3.1-SNAP-3xHA or pBabe-H3.3-SNAP-3xHA and selection in medium containing 5 µg/ml blasticidin. Cells were then infected with the indicated shRNA, selected with puromycin for 48 hr, and seeded in Lab-Tek II Chamber slides (Labclinics). After 48 hr, SNAP labelling was performed as previously described (Bodor et al., 2012), with modifications in compound concentration and treatment time.

For specific labelling of newly-synthetized histones (quench-chase-pulse experiments), pre-existing histones were first quenched by incubating cells with 5 µM SNAP-cell Block (New England Biolabs; S9106S) for 30 min at 37 °C. After two washes with medium, cells were incubated in medium for 30 min, washed again twice, and incubated in fresh medium at 37°C for the chase period (6 to 7 hr). The pulse step was performed with 2 µM SNAP-cell TMR Star (New England Biolabs, S9105S) for 15 min followed by two washes with medium, incubation with fresh medium for 30 min and two more washes. At this point, cells were pre-extracted in 0.2% Triton X-100/PBS for 5 min on ice to remove the unbound chromatin fraction and fixed in 4% paraformaldehyde for 10 min at room temperature. Fixed cells were permeabilized with 0.2% Triton X-100/PBS for 10 min at room temperature and stained with PBS-DAPI (0.25 µg/ml).

High-throughput microscopy (HTM)-mediated quantification of the nuclear intensity of SNAP-H3.1/3 was performed as described (Lee et al., 2018). Forty-eight images per well were automatically acquired with a robotized fluorescence microscopy station (ScanR, Olympus) at 40× magnification. Signals were calculated with CellProfiler using DAPI staining to generate masks of cell nuclei. Uneven background from fluorescence microscopy images and artifacts from autofluorescence were removed with ImageJ to ensure a proper quantification.

### Clonogenic Assay

Cells were plated into a 6-well plate at a clonogenic density (500, 750, or 1,000 cells/well) with puromycin selection, replacing the media every 3 days. After 7 to 15 days, media was removed, and cells were washed with PBS and stained with a mixture of 6% glutaraldehyde and 0.5% crystal violet for at least 30 min. Plates were then washed by careful immersion into water and air-dried at room temperature.

### MTT Assay for Cell Proliferation

After infection and selection for 48 hr with puromycin, 10,000 cells/well were seeded in 96-well plates with triplicates. The following days, cells were incubated with MTT (3-(4,5-dimethylthiazol-2-yl)-2,5-diphenyltetrazolium bromide; Sigma) (0.5 mg/ml) in Dulbecco’s modified Eagle’s medium without FBS for 3 hr at 37 °C. After incubation, dimethyl sulfoxide-isopropanol (1:4) was added; once the purple crystals were fully dissolved, the absorbance was measured at 570 nm.

### Migration and Invasion Assays

For migration assays, 30,000 cells were resuspended in DMEM 0.1% FBS and 0.1% BSA, and reseeded on a transwell filter chamber (Corning; 3422). After 1 hr, DMEM 10% FBS was added to the lower chamber as a chemoattractant and incubated for 6 to 8 hr. For invasion experiments, cells were placed in Matrigel-coated transwell (Corning; 354230) and incubated for 12 to 16 hr after chemoattractant addition.

Non-migrating and non-invading cells were removed from the upper surface of the membrane, whereas cells adhered to the lower surface were fixed with PFA 4% for 15 min and nuclei-stained with PBS-DAPI (0.25 µg/ml). Images were acquired with InCell 2000 automated epifluorescence microscope, and DAPI-stained nuclei were counted using ImageJ software.

### Rescue Experiments with Suv39H1-GFP

For Suv39H1-GFP rescue experiments, MDA-MB-231 cells were infected with the indicated shRNAs and selected with puromycin for 48 hr. Cells were then seeded and infected twice with the retroviral particles expressing pBabe-Suv39H1-GFP or pBabe-GFP (as described above). Migration experiments were performed at 48 hr after the second infection, and GFP-positive cells were counted using the ImageJ software.

### Tumor Xenografts

Mice were maintained and treated in accordance with institutional guidelines of Vall d’Hebron University Hospital Care and Use Committee. MDA-MB-231 cells were infected with shControl, shLOXL2, shRBX1, shRUVBL2, shH2A.Z.1 or shDDB1 and selected for 48 hr with puromycin. 48 hr post-selection, 1 × 10^6^ cells were injected into the mammary fat pad of gland number 4 of 6-week-old female BALB/c nude mice purchased from Charles Rivers Laboratories (L’Arbresle Cedex, France). Tumour xenografts were measured using a calliper twice per week, and tumour volume was calculated using the formula (length x width^2^) x (pi/6). Mice were sacrificed by cervical dislocation at the end of the experiment.

### Cloning Procedures and Plasmids

pBabe-GFP or pBabe-SUV39H1 were generated by subcloning CGA-pCAGGS-Suv39H1-EGFP-IRES-Puro vector into the pBabe-Puro expression vector (Addgene #1764). Suv39H1-GFP or ATG-GFP sequences were first amplified by PCR from the CGA-pCAGGS-Suv39H1-EGFP-IRES-Puro vector. The pBabe target vector was digested with BamHI and SalI, and the insert sequences were introduced using Gibson assembly.

### Statistical Analysis

Statistical significance was assessed using an unpaired, two-tailed Student’s t-test. For in vivo experiments, two-way ANOVA with subsequent Bonferroni correction and Log-rank (Mantel-Cox) test were used to assess statistical significance in tumor growth and tumor incidence, respectively.

### Accession Numbers

Raw data and processed information of the ChIPseq, RNA-seq, and ATAC-seq experiments generated in this article were deposited in the National Center for Biotechnology Information Gene Expression Omnibus (NCBI GEO) (Barrett et al., 2013) repository under the accession number GSE142463.

## RESOURCES TABLE

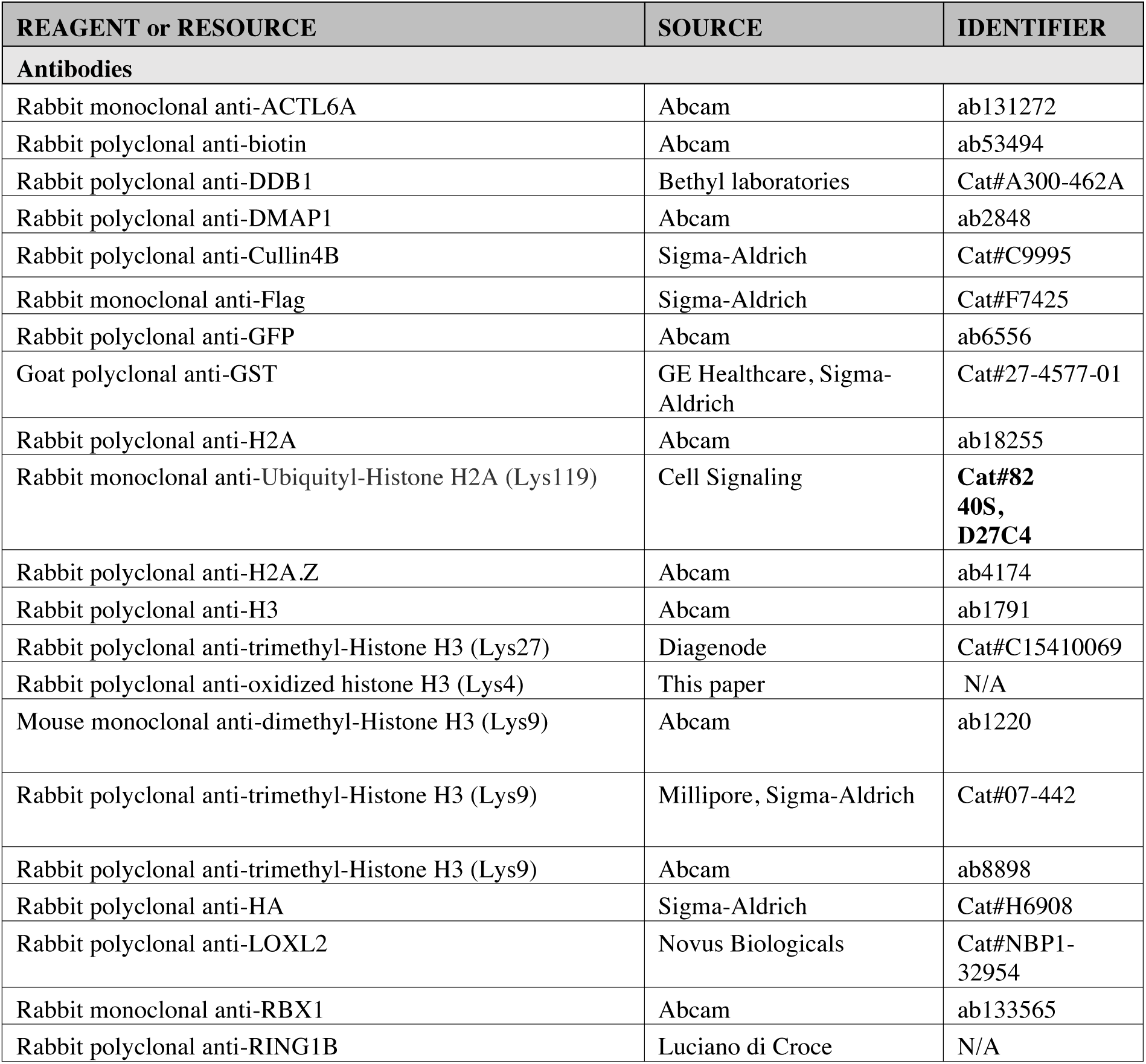
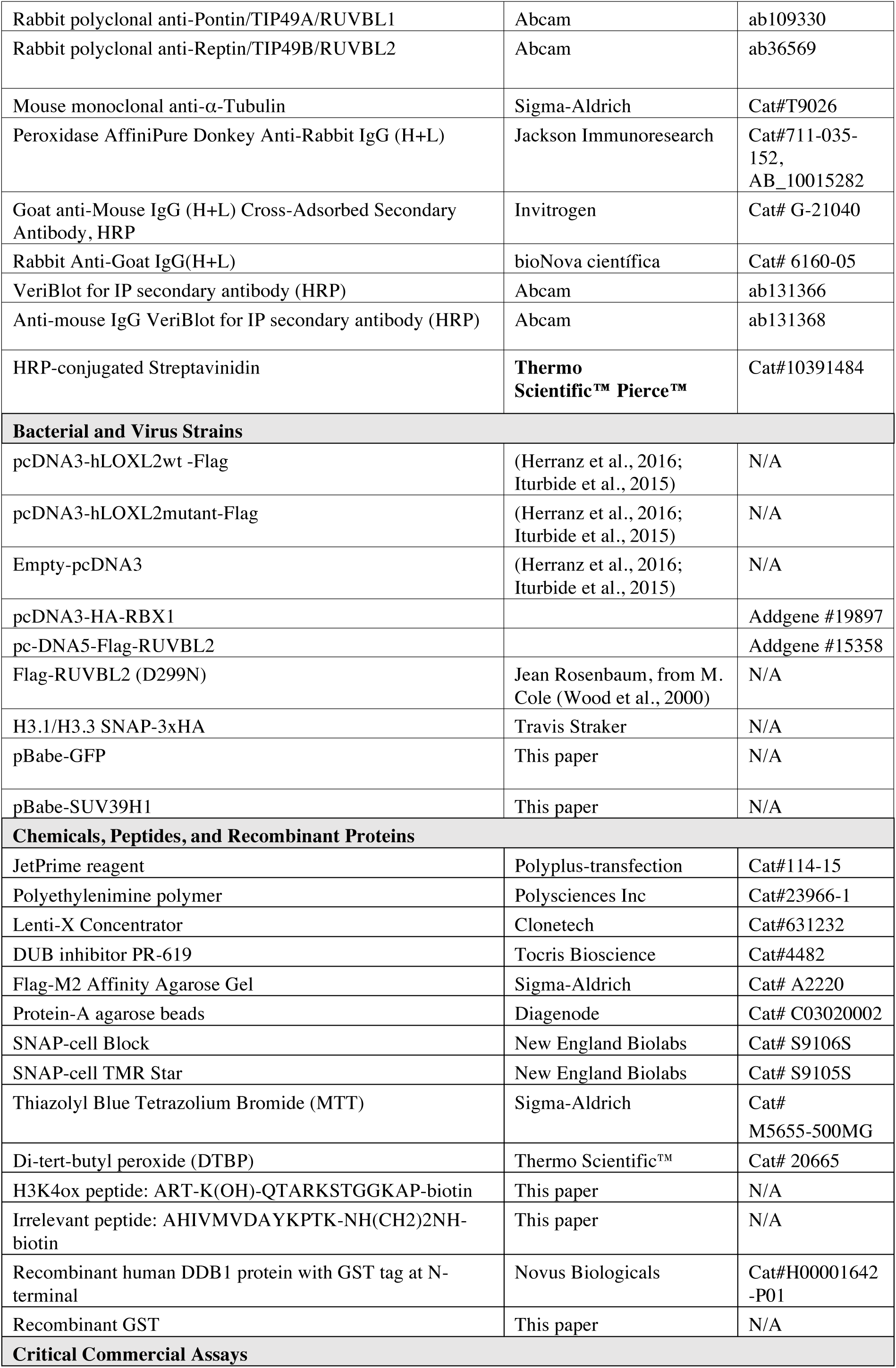
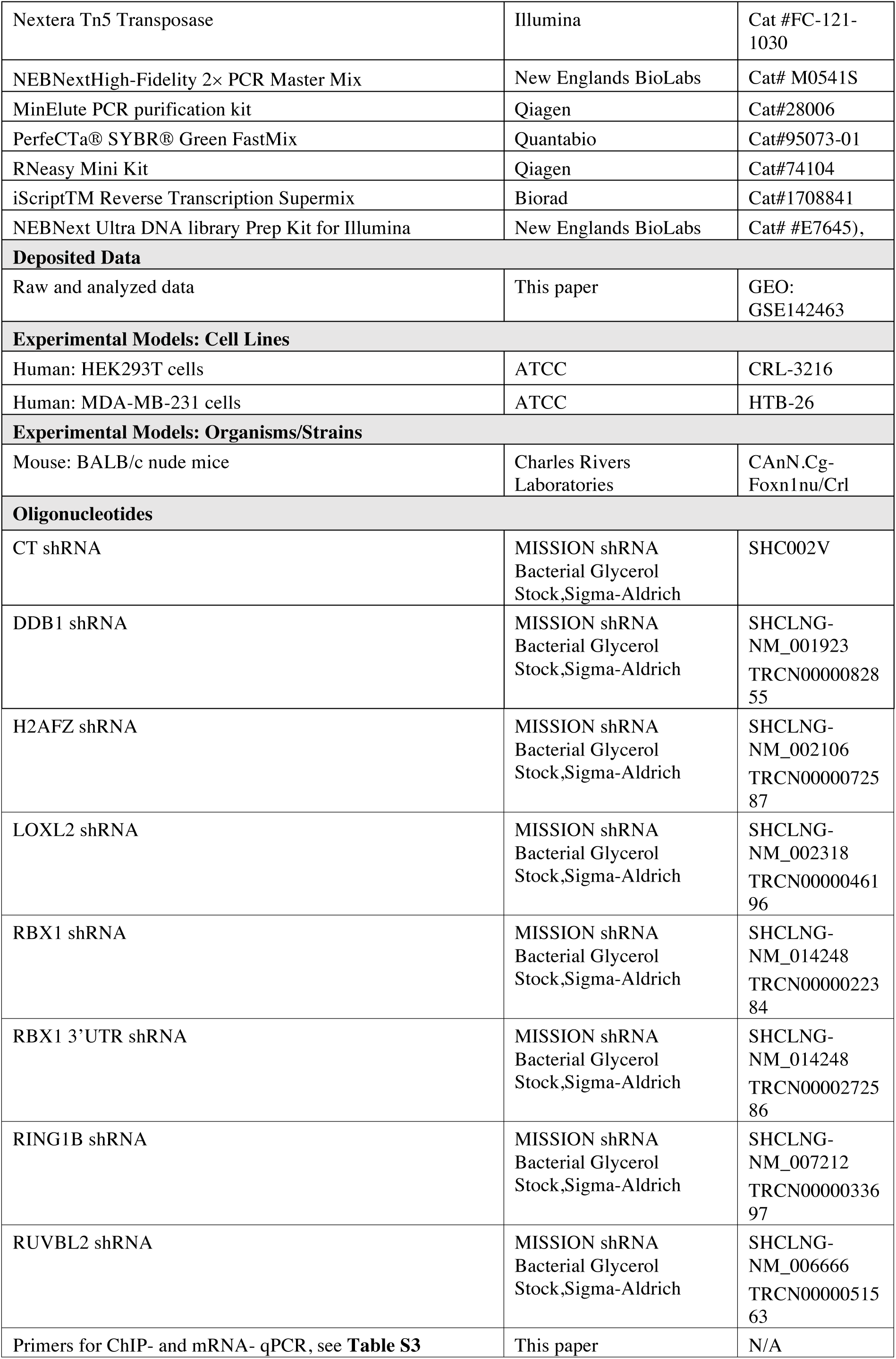
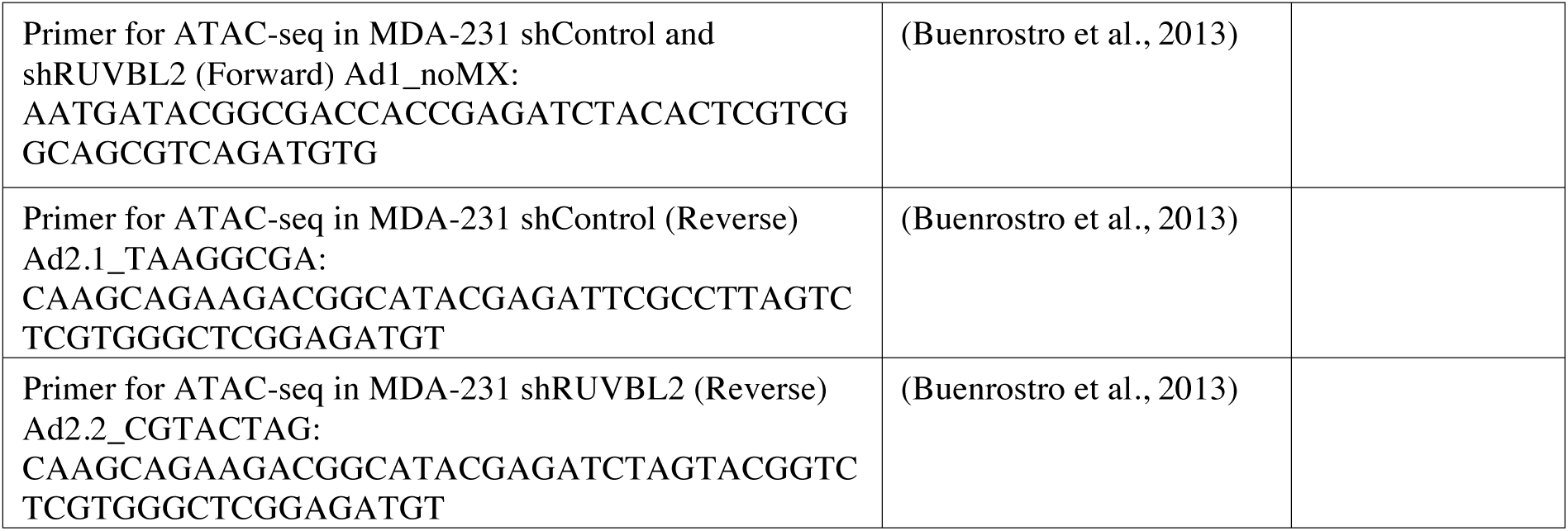

