## Supplemental information for "Maintaining oxidized H3 in heterochromatin is required for the oncogenic capacity of triple-negative breast cancer cells"

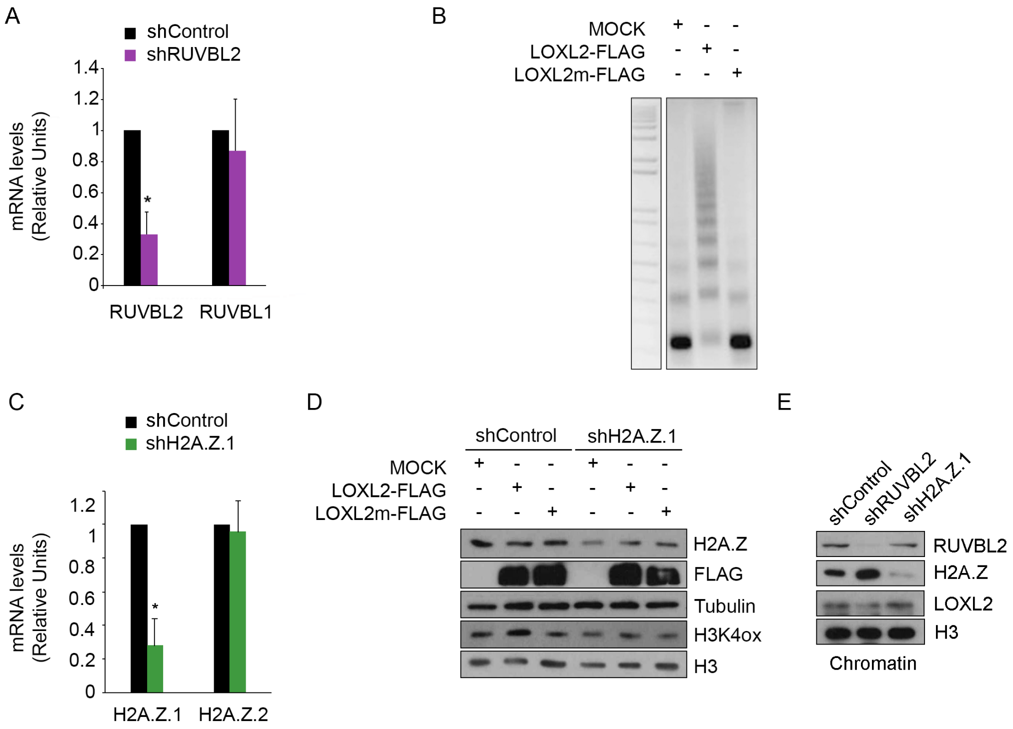


**Figure S1, related to Figure 1. RUVBL2 and H2A.Z are required for LOXL2 induction of chromatin compaction.**

**A)** mRNA levels of RUVBL1 and RUVBL2 analyzed by qRT-PCR in HEK293T cells infected with shControl or shRUVBL2. Gene expression was normalized to the *Pumilio* housekeeping gene and is presented as the fold-change relative to the shControl cells, which was set as 1. Error bars indicate standard deviation from at least three experiments. $*p < 0.05$.

**B)** HEK293T cells were transfected with wild-type LOXL2, a catalytically-inactive LOXL2 mutant (LOXL2m), or an empty vector (mock). At 48 hr post-transfection, isolated nuclei were digested with micrococcal nuclease (MNase) for 2 min, and total genomic DNA was analyzed using agarose gel electrophoresis. Intervening lanes were removed as indicated.

**C)** mRNA levels of H2A.Z.1 and H2A.Z.2 analyzed by qRT-PCR in HEK293T cells infected with shControl or shH2A.Z.1. Gene expression was normalized to the *Pumilio* housekeeping gene and presented as the fold-change relative to the shControl cells, which was set as 1. Error bars indicate standard deviation in at least three experiments. *$p < 0.05$.

**D)** HEK293T cells infected with shControl or shH2A.Z.1 were transfected with wild-type LOXL2, the inactive mutant LOXL2m, or an empty vector (mock). At 48 hr after transfection, total and histone extracts were obtained and analyzed by Western blot with the indicated antibodies.

**E)** Subcellular fractionation assays were performed in MDA-MB-231 cells infected with shControl, shRUVBL2, or shH2A.Z.1. The chromatin fraction was analyzed by Western blot with the indicated antibodies.

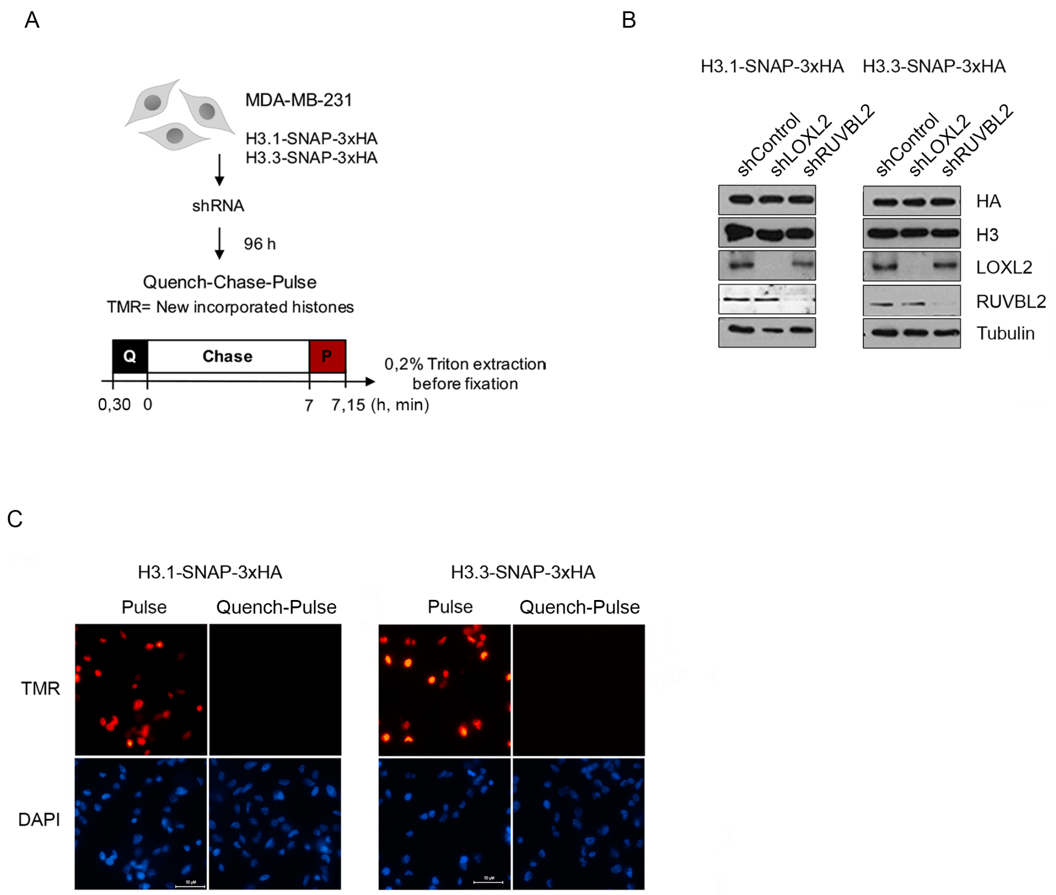


**Figure S2, related to Figure 2. *In vivo* Visualization Assay for Newly Synthesized H3.1 and H3.3.**

**A)** Experimental design for specific labeling of newly-synthetized histones H3.1 and H3.3. MDA-MB-231 cells stably expressing H3.1-SNAP or H3.3-SNAP were infected with shControl, shLOXL2, or shRUVBL2. After 96 hr, quench-chase-pulse experiments were performed: cells were quenched with 5 µM SNAP-cell block for 30 min at 37ºC, washed, and incubated in fresh medium at 37ºC for the chase period (7 hr). The pulse step was performed with 2 µM SNAP-cell TMR Star for 15 min. Cells were pre-extracted in 0.2% Triton X-100/PBS for 5 min on ice to remove the unbound chromatin fraction and fixed in 4% paraformaldehyde for 10 min at room temperature.

**B)** Total extracts of MDA-MB-231 cells stably expressing H3.1/3-SNAP at 96 hr after shRNA infection. Protein levels were analyzed by Western blot using the indicated antibodies.

**C)** Fluorescent microscopy visualization of MDA-MB-231 cells expressing H3.1/H3.3-SNAP after labeling assays with red fluorescent TMR-Star. The TMR pulse labels pre-existing H3.1/3-SNAP, and the quench control ensures that treatment with a non-fluorescent block prevents further labeling with TMR-Star. Nuclei were stained with DAPI. Scale bars represent 50 μM.

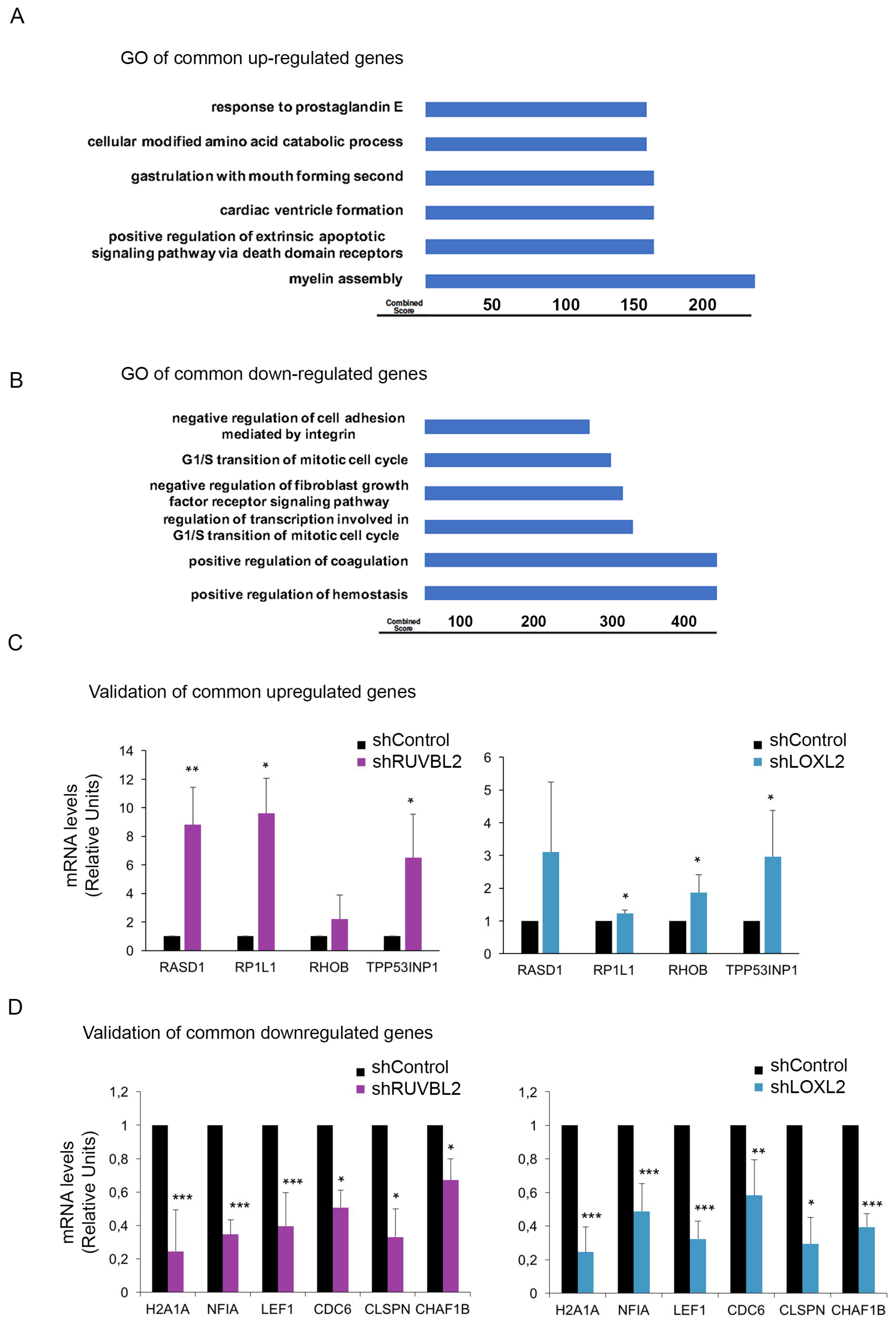


**Figure S3, related to Figure 3. Common Dysregulated Genes in shRUVBL2 and shLOXL2 knockdown cells.**

**A, B)** Gene ontology analysis (biological process 2018) of commonly upregulated (A) and the downregulated (B) genes in MDA-MB-231 cells infected with shControl, shLOXL2, or shRUVBL2**.**

**C, D)** Validation of the RNA-sequencing results. MDA-MB-231 cells were infected with shControl, shLOXL2, or shRUVBL2, and mRNA levels of commonly upregulated (C) or downregulated (D) genes were analyzed by qRT-PCR. Gene expression was normalized to the *Pumilio* housekeeping gene and are presented as the fold-change relative to the shControl cells, which was set as 1. Error bars indicate standard deviation in at least three experiments. $^{*}p < 0.05$, $^{**}p \leq 0.01$, $^{***}p \leq 0.001$.

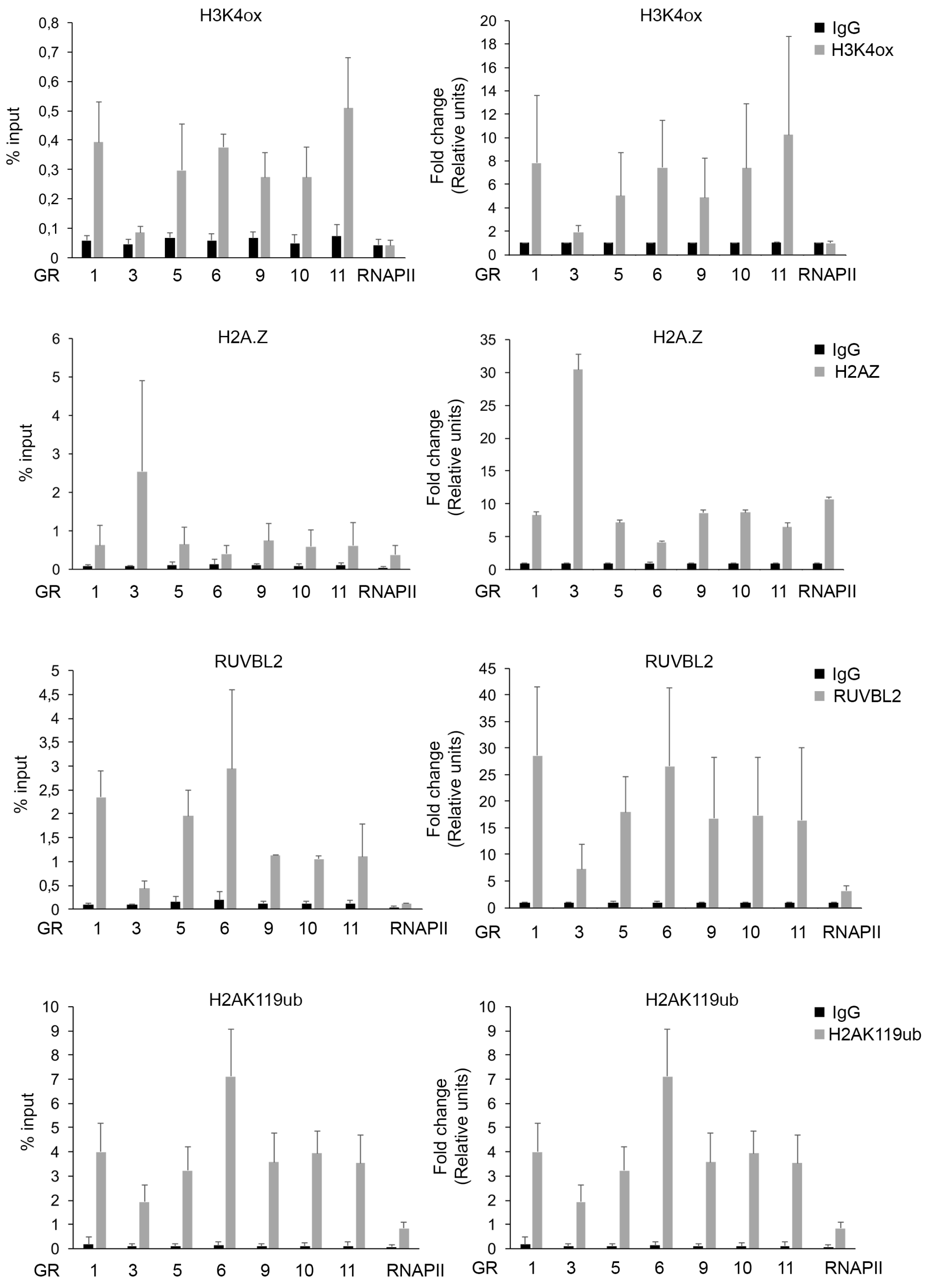


**Figure S4, related to Figure 4 and Figure 6. ChIPseq validation.**

ChIP-PCR experiments of H3K4ox, RUVBL2, H2A.Z and H2AK119ub were performed in selected genomic regions (GR) and at the RNAPII promoter (RNAPII), which was used as a negative control for H3K4ox, RUVBL2 and H2AK119ub. Data of qPCR amplifications were normalized to the input and to total immunoprecipitated H3 or H2A for H3K4ox and

H2AK119ub, respectively. Results are expressed as percentage of input (left) or fold change (right) relative to IgG binding. Error bars indicate SD in at least three experiments. $*p < 0.05$, $**p \leq 0.01$, $***p \leq 0.001$.

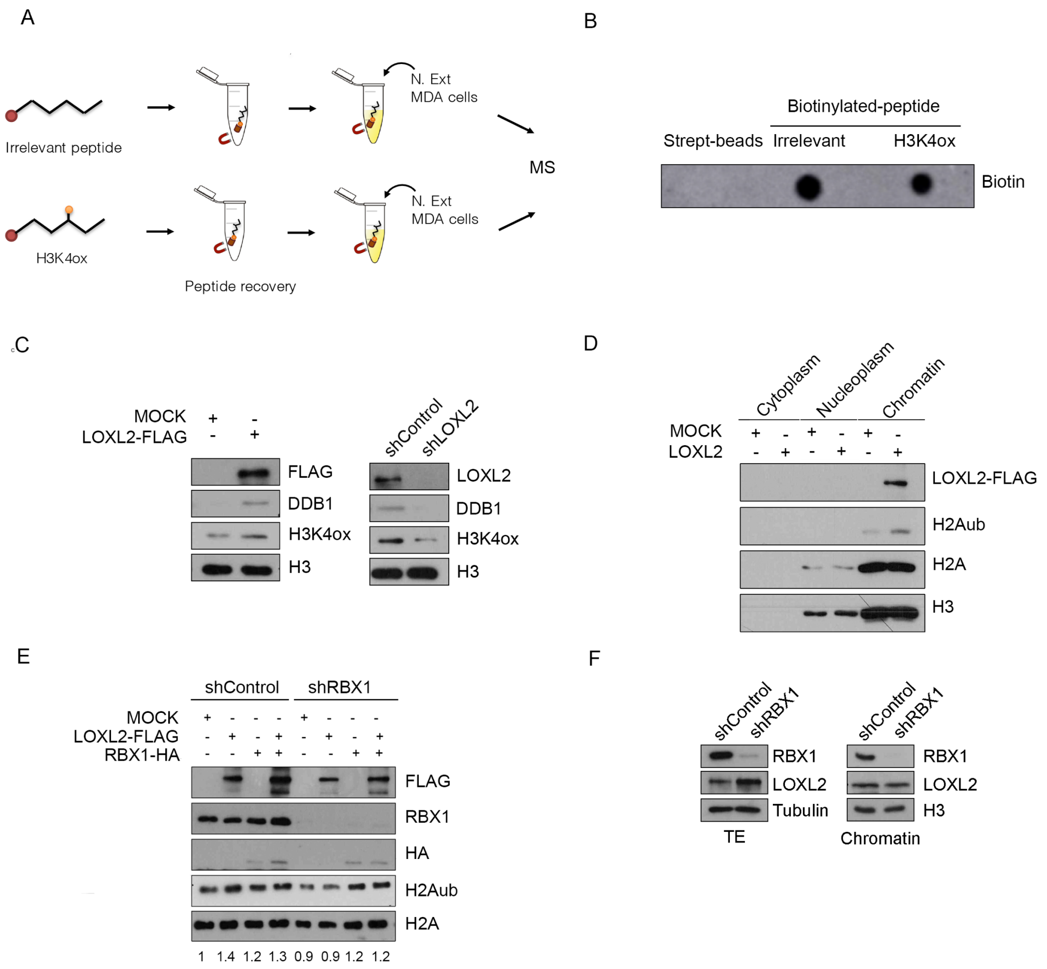


**Figure S5, related to Figure 5. DDB1 is a H3K4ox reader involved in the ubiquitination of histone H2A through the CRL4B complex.**

**A)** Schematic representation of the experimental approach used to identify putative H3K4ox readers. Peptides were immobilized with streptavidin beads, washed, and incubated with MDA-MB-231 nuclear extracts. Bound proteins were subjected to trypsin digestion and analyzed by liquid chromatography–tandem mass spectrometry (LC-MS/MS).

**B)** Recovery of the biotinylated peptides was analyzed by dot blot with an anti-biotin antibody.

**C)** HEK293T cells were transfected with an empty vector or LOXL2-FLAG, and cell extracts were obtained 48 hr later (left panel). MDA-MB-231 cells were infected with shControl or shLOXL2 and selected for 96 hr (right panel). Subcellular fractionation assays were performed, and the chromatin fraction was analyzed by Western blot with the indicated antibodies.

**D)** HEK293T cells were transfected with LOXL2-FLAG or an empty vector. After 48 hr, the cell extracts were fractionated. Protein levels were analyzed by Western blot with the indicated antibodies.

**E)** HEK293T cells infected with shControl or shRBX1 were transfected with empty vector, LOXL2-FLAG or RBX1-HA, and chromatin association assays were performed after 48 hr. Protein levels were analyzed by Western blot using the indicated antibodies. Numbers indicate quantification of H2Aub normalized to the total levels of H2A; values are presented as the fold-change relative to the shControl cells transfected to mock vector, which was set to 1.

**F)** MDA-MB-231 cells were infected with shControl or shRBX1. Total extracts (left panel) and the chromatin fraction obtained by subcellular fractionation assays (right panel) were analyzed by Western blot with the indicated antibodies.

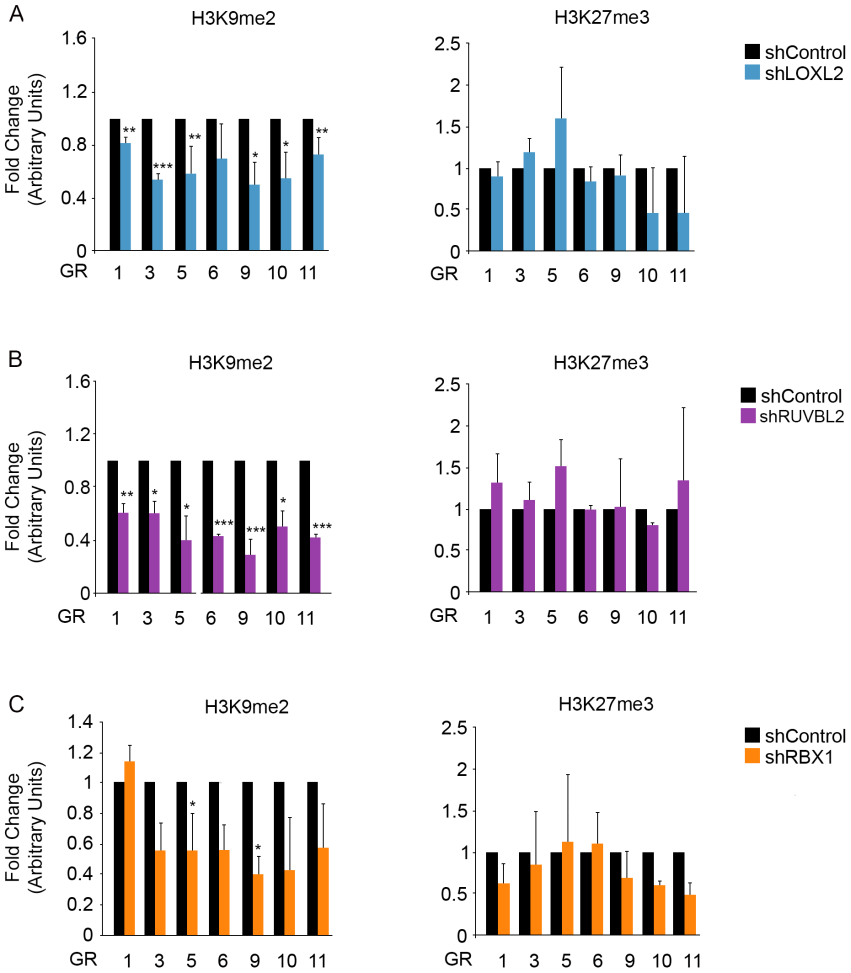


**Figure S6, related to Figure 7. LOXL2, RUVBL2, and RBX1 Regulate Levels of H3K9me2 but Not of H3K27me3 in MDA-MB-231 Cells.**

**A–C)** MDA-MB-231 cells were infected with shControl, shLOXL2 **(A),** shRUVBL2 **(B),** or shRBX1 **(C)**. ChIP-PCR experiments of H3K9me2 and H3K27me3 were performed in the selected genomic regions (GR). Data of qPCR amplifications were normalized to the input and to total immunoprecipitated H3. Results are expressed as fold-changes relative to the data obtained in shControl, which was set to 1. Error bars indicate SD from at least three experiments. $*p < 0.05$, $**p \leq 0.01$, $***p \leq 0.001$.

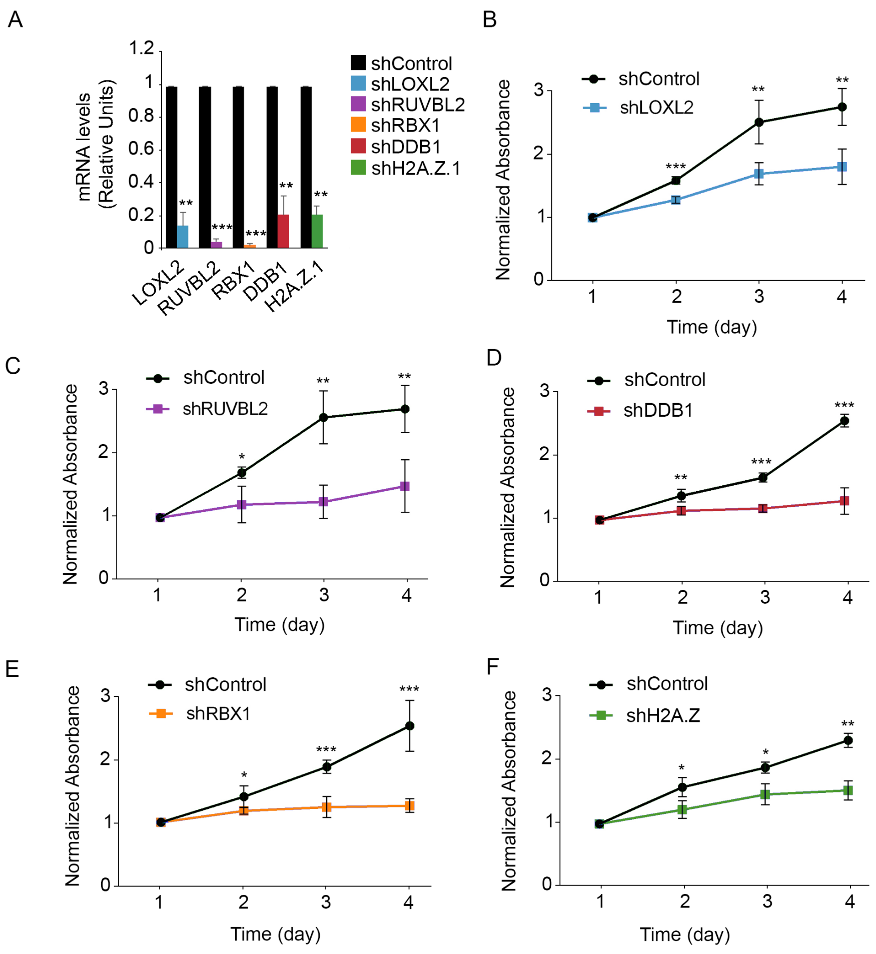


**Figure S7, related to Figure 8. Knockdown of LOXL2, RUVBL2, DDB1, RBX1, or H2A.Z.1 Decreases the Proliferation Rates of MDA-MB-231 Cells.**

**A)** MDA-MB-231 cells were infected with shControl, shLOXL2, shRUVBL2, shDDB1, shRBX1 or shH2A.Z.1. mRNA levels of the indicated genes were analyzed by qRT-PCR. Gene expression levels were normalized to the *Pumilio* housekeeping gene and are presented as fold-change relative to the shControl cells.

**B–F)** MTT assays in MDA-MB-231 cells infected with shControl or shLOXL2 (B), shRUVBL2 (C), shDDB1 (D), shRBX1 (E), or shH2A.Z.1 (F). Measurements were obtained over four consecutive days after selection. Error bars indicate the SD from at least three independent experiments. *$p < 0.05$, **$p \leq 0.01$, ***$p \leq 0.001$.

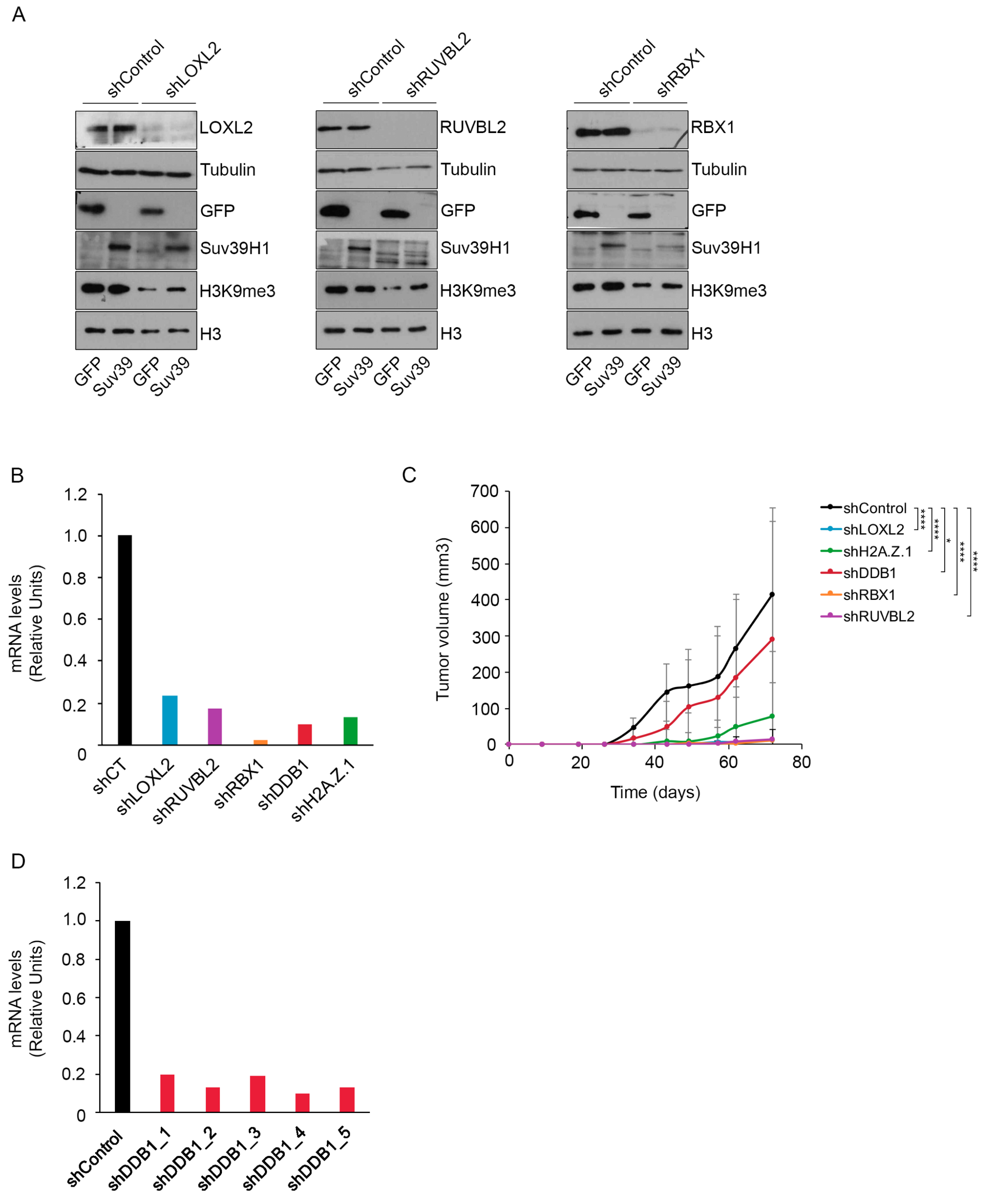


**Figure S8, related to Figure 8. Heterochromatin alterations blocks the oncogenic properties of MDA-MB-231 breast cancer cells.**

**A)** MDA-MB-231 cells were first infected with shControl, shLOXL2, shRUVBL2 or shRBX1. After puromycin selection, knocked-down cells were reinfected with GFP or SUV39H1-GFP and total cell extracts were obtained 48 hr later. Protein levels were analyzed by Western blot using the indicated antibodies.

**B)** MDA-MB-231 cells were infected with shControl, shLOXL2, shRUVBL2, shDDB1, shRBX1 or shH2A.Z.1. Before injecting cells into the mice, knockdown efficiency was checked by analysing the mRNA levels of the indicated genes by qRT-PCR. Gene expression levels were normalized to the *HPRT* housekeeping gene and presented as fold-change relative to the shControl cells.

**C)** MDA-MB-231 cells were infected with shControl, shLOXL2, shDDB1, shH2A.Z.1, shRUVBL2, or shRBX1. After puromycin selection, 1x106 knocked-down cells were injected into the mammary fat pad of nude mice. Tumor volume was monitored at the indicated time points and the results were expressed as averages ± S.D. Indicated statistical analysis was performed with values at day 72; in all cases, knocked-down groups were compared with shControl $*p < 0.05$, $**p \leq 0.01$, $***p \leq 0.001$.

**D)** At the end of the experiment, RNA was extracted from tumor samples and mRNA levels of DDB1 were analysed by qRT-PCR. Gene expression levels were normalized to the *HPRT* housekeeping gene and presented as fold-change relative to the shControl cells.

| GENE | MS/MS SCORE |
|---|---|
| **IPI00294839 Tax_Id=9606 Gene_Symbol=LOXL2;ENTPD4 Lysyl oxidase homolog 2** | **1934** |
| IPI00003362 Tax_Id=9606 Gene_Symbol=HSPA5 HSPA5 protein | 1561 |
| IPI00304925 Tax_Id=9606 Gene_Symbol=HSPA1A;HSPA1B Heat shock 70 kDa protein 1 | 1439 |
| IPI00396485 Tax_Id=9606 Gene_Symbol=EEF1A1 Elongation factor 1-alpha 1 | 868 |
| IPI00003865 Tax_Id=9606 Gene_Symbol=HSPA8 Isoform 1 of Heat shock cognate 71 kDa protein | 559 |
| **IPI00021187 Tax_Id=9606 Gene_Symbol=RUVBL1 Isoform 1 of RuvB-like 1** | **411** |
| IPI00397801 Tax_Id=9606 Gene_Symbol=FLG2 Filaggrin-2 | 305 |
| **IPI00009104 Tax_Id=9606 Gene_Symbol=RUVBL2 RuvB-like 2** | **257** |
| IPI00221127 Tax_Id=9606 Gene_Symbol=MYLK2 Myosin light chain kinase 2, skeletal/cardiac muscle | 226 |
| IPI00012837 Tax_Id=9606 Gene_Symbol=KIF5B Kinesin-1 heavy chain | 204 |
| **IPI00003935 Tax_Id=9606 Gene_Symbol=HIST2H2BE Histone H2B type 2-E** | **155** |
| IPI00386597 Tax_Id=9606 Gene_Symbol=SPRR2E Small proline-rich protein 2E | 150 |
| IPI00179330 Tax_Id=9606 Gene_Symbol=UBB;RPS27A;UBC ubiquitin and ribosomal protein S27a precursor | 150 |
| IPI00183572 Tax_Id=9606 Gene_Symbol=DOCK7 Isoform 2 of Dedicator of cytokinesis protein 7 | 108 |
| IPI00395474 Tax_Id=9606 Gene_Symbol=SAMD1 Atherin | 98 |
| IPI00083708 Tax_Id=9606 Gene_Symbol=BAT2D1 Isoform 7 of BAT2 domain-containing protein 1 | 90 |
| **IPI00219919 Tax_Id=9606 Gene_Symbol=DMAP1 DNA methyltransferase 1-associated protein 1** | **75** |
| IPI00017297 Tax_Id=9606 Gene_Symbol=MATR3 Matrin-3 | 74 |
| IPI00022204 Tax_Id=9606 Gene_Symbol=SERPINB3 Isoform 1 of Serpin B3 | 73 |
| IPI00182106 Tax_Id=9606 Gene_Symbol=MEN1 Isoform 2 of Menin | 71 |
| IPI00876962 Tax_Id=9606 Gene_Symbol=INF2 Isoform 2 of Inverted formin-2 | 64 |
| IPI00021634 Tax_Id=9606 Gene_Symbol=KLC2 Kinesin light chain 2 | 59 |
| IPI00017669 Tax_Id=9606 Gene_Symbol=SMARCE1 Isoform 1 of SWI/SNF-related matrix-associated actin-dependent regulator of chromatin subfamily E member 1 | 58 |
| IPI00304903 Tax_Id=9606 Gene_Symbol=SPRR1B Cornifin-B | 57 |
| IPI00217652 Tax_Id=9606 Gene_Symbol=GLT8D3 Isoform 1 of Glycosyltransferase 8 domain-containing protein 3 | 56 |
| IPI00008603 Tax_Id=9606 Gene_Symbol=ACTA2 Actin, aortic smooth muscle | 55 |
| IPI00172636 Tax_Id=9606 Gene_Symbol=CAMK2D Isoform Delta 6 of Calcium/calmodulin-dependent protein kinase type II delta chain | 53 |
| **IPI00171611 Tax_Id=9606 Gene_Symbol=HIST2H3A;HIST2H3C;HIST2H3D Histone H3.2** | **49** |

| | |
|---|---|
| IPI00022462 Tax_Id=9606 Gene_Symbol=TFRC Transferrin receptor protein 1 | 47 |
| IPI00332371 Tax_Id=9606 Gene_Symbol=PFKL Isoform 1 of 6-phosphofructokinase, liver type | 47 |
| IPI00101659 Tax_Id=9606 Gene_Symbol=NSUN5 Isoform 2 of Putative methyltransferase NSUN5 | 45 |
| IPI00027547 Tax_Id=9606 Gene_Symbol=DCD Dermcidin | 45 |
| IPI00465431 Tax_Id=9606 Gene_Symbol=LGALS3 Galectin-3 | 44 |
| IPI00328987 Tax_Id=9606 Gene_Symbol=BYSL Bystin | 40 |
| IPI00179964 Tax_Id=9606 Gene_Symbol=PTBP1 Isoform 1 of Polypyrimidine tract-binding protein 1 | 38 |
| **IPI00003627 Tax_Id=9606 Gene_Symbol=ACTL6A Isoform 1 of Actin-like protein 6A** | **36** |
| IPI00013885 Tax_Id=9606 Gene_Symbol=CASP14 Caspase-14 | 36 |
| IPI00290950 Tax_Id=9606 Gene_Symbol=WDR20 Isoform 2 of WD repeat-containing protein 20 | 35 |
| IPI00081836 Tax_Id=9606 Gene_Symbol=HIST1H2AH Histone H2A type 1-H | 34 |
| IPI00410330 Tax_Id=9606 Gene_Symbol=GATAD2A Isoform 1 of Transcriptional repressor p66-alpha | 33 |
| IPI00172450 Tax_Id=9606 Gene_Symbol=CAMK2G Isoform 4 of Calcium/calmodulin-dependent protein kinase type II gamma chain | 32 |
| IPI00031008 Tax_Id=9606 Gene_Symbol=TNC Isoform 1 of Tenascin | 30 |
| IPI00290566 Tax_Id=9606 Gene_Symbol=TCP1 T-complex protein 1 subunit alpha | 30 |
| IPI00022200 Tax_Id=9606 Gene_Symbol=COL6A3 Isoform 1 of Collagen alpha-3(VI) chain | 29 |
| IPI00025753 Tax_Id=9606 Gene_Symbol=DSG1 Desmoglein-1 | 29 |
| IPI00334579 Tax_Id=9606 Gene_Symbol=MRPL43 mitochondrial ribosomal protein L43 isoform d | 28 |
| IPI00457109 Tax_Id=9606 Gene_Symbol=ABCA12 Isoform 2 of ATP-binding cassette sub-family A member 12 | 28 |
| IPI00298860 Tax_Id=9606 Gene_Symbol=LTF Growth-inhibiting protein 12 | 28 |
| IPI00006158 Tax_Id=9606 Gene_Symbol=LRMP Lymphoid-restricted membrane protein | 27 |
| IPI00299554 Tax_Id=9606 Gene_Symbol=KIF14 Kinesin-like protein KIF14 | 27 |
| IPI00643923 Tax_Id=9606 Gene_Symbol=BAT2L KIAA0515 | 26 |
| IPI00101532 Tax_Id=9606 Gene_Symbol=CEP55 Isoform 1 of Centrosomal protein of 55 kDa | 26 |
| IPI00017617 Tax_Id=9606 Gene_Symbol=DDX5 Probable ATP-dependent RNA helicase DDX5 | 26 |
| IPI00168899 Tax_Id=9606 Gene_Symbol=DBF4B Isoform 1 of Protein DBF4 homolog B | 26 |
| IPI00298202 Tax_Id=9606 Gene_Symbol=ACOT8 Acyl-coenzyme A thioesterase 8 | 25 |
| IPI00031552 Tax_Id=9606 Gene_Symbol=ELAVL3 Isoform 1 of ELAV-like protein 3 | 24 |

| | |
|---|---|
| IPI00395573 Tax_Id=9606 Gene_Symbol=OPRM1 Opioid receptor, mu 1 | 24 |
| IPI00247674 Tax_Id=9606 Gene_Symbol=PALM3 Paralemmin-3 | 23 |
| IPI00003505 Tax_Id=9606 Gene_Symbol=TRIP13 Isoform 1 of Thyroid receptor-interacting protein 13 | 23 |

**Table S1, related to Figure 1. New list of putative LOXL2 interactors**. List of putative LOXL2 interactors identified in a tandem-affinity purification approach and mass spectrometry analysis. Gene symbols and MS scores are shown.

| | Prot.Nm | T | C | LogFC | p.value | adjp | DEP |
|---|---|---|---|---|---|---|---|
| NOL11_HUMAN | NOL11 | 12,3 | 0 | 37 | 3,17E-05 | 3,67E-03 | TRUE |
| NUCL_HUMAN | NCL | 8 | 0 | 36,17 | 3,61E-02 | 0,0007327 | TRUE |
| HNRL1_HUMAN | HNRNPUL1 | 5,7 | 0 | 35,82 | 0,0002448 | 0,003614 | TRUE |
| NIP7_HUMAN | NIP7 | 5 | 0 | 35,77 | 0,000268 | 0,003818 | TRUE |
| BAF_HUMAN | BANF1 | 4,3 | 0 | 35,69 | 0,0004804 | 0,006142 | TRUE |
| RRP7A_HUMAN | RRP7A | 5,7 | 0 | 35,46 | 0,0008638 | 0,009742 | TRUE |
| LACTB_HUMAN | LACTB | 4,3 | 0 | 35,37 | 0,001791 | 0,01634 | TRUE |
| DDB1_HUMAN | DDB1 | 3,3 | 0 | 35,32 | 0,001876 | 0,01692 | TRUE |
| PNO1_HUMAN | PNO1 | 4 | 0 | 35,17 | 0,001633 | 0,01542 | TRUE |
| PK1IP_HUMAN | PAK1IP1 | 3,7 | 0 | 35,03 | 0,00374 | 0,02839 | TRUE |
| SRFB1_HUMAN | SRFBP1 | 3,7 | 0 | 34,95 | 0,006841 | 0,04553 | TRUE |
| RBM19_HUMAN | RBM19 | 3,7 | 0 | 34,95 | 0,006841 | 0,04553 | TRUE |
| AKAP8_HUMAN | AKAP8 | 3,7 | 0 | 34,87 | 0,006525 | 0,04452 | TRUE |
| EXOS6_HUMAN | EXOSC6 | 3 | 0 | 34,87 | 0,007173 | 0,04697 | TRUE |
| RRP36_HUMAN | RRP36 | 4 | 0 | 34,87 | 0,007173 | 0,04697 | TRUE |
| UTP15_HUMAN | UTP15 | 15,3 | 0,5 | 4,715 | 3,02E-05 | 3,67E-03 | TRUE |
| RL35A_HUMAN | RPL35A | 9,3 | 0,5 | 3,989 | 4,35E-02 | 0,000861 | TRUE |
| NLE1_HUMAN | NLE1 | 6,7 | 0,5 | 3,56 | 0,0007656 | 0,008986 | TRUE |
| PHIP_HUMAN | PHIP | 15 | 1 | 3,547 | 2,18E-03 | 0,0001265 | TRUE |
| RLA0_HUMAN | RPLP0 | 14,3 | 1 | 3,501 | 3,62E-03 | 0,0001727 | TRUE |
| XRN2_HUMAN | XRN2 | 12,3 | 1 | 3,385 | 1,15E-02 | 0,0003448 | TRUE |
| REXO4_HUMAN | REXO4 | 12 | 1 | 3,333 | 1,90E-02 | 0,0004811 | TRUE |
| DDX31_HUMAN | DDX31 | 5,3 | 0,5 | 3,318 | 0,002642 | 0,02227 | TRUE |
| RS5_HUMAN | RPS5 | 12 | 1 | 3,274 | 3,24E-02 | 0,0006928 | TRUE |
| RLA0L_HUMAN | RPLP0P6 | 11,3 | 1 | 3,264 | 3,49E-05 | 0,0007267 | TRUE |
| DHX30_HUMAN | DHX30 | 10 | 1 | 3,155 | 9,09E-02 | 0,001639 | TRUE |
| NEP1_HUMAN | EMG1 | 5,3 | 0,5 | 3,088 | 0,00746 | 0,04808 | TRUE |
| WDR74_HUMAN | WDR74 | 19,3 | 2 | 3,064 | 1,16E-04 | 8,55E-03 | TRUE |
| PWP2_HUMAN | PWP2 | 14,7 | 1,5 | 3,064 | 4,44E-03 | 0,0001801 | TRUE |
| HEAT1_HUMAN | HEATR1 | 20 | 2,5 | 2,959 | 2,07E-05 | 3,35E-03 | TRUE |
| RL9_HUMAN | RPL9 | 9 | 1 | 2,918 | 0,0005109 | 0,00621 | TRUE |
| WDR3_HUMAN | WDR3 | 23,7 | 3 | 2,681 | 1,12E-04 | 8,55E-03 | TRUE |
| TBL3_HUMAN | TBL3 | 24,7 | 4 | 2,535 | 1,72E-05 | 3,35E-03 | TRUE |
| DDX52_HUMAN | DDX52 | 5 | 1 | 2,444 | 0,007399 | 0,04807 | TRUE |
| EXOSX_HUMAN | EXOSC10 | 10 | 1,5 | 2,399 | 0,001388 | 0,01342 | TRUE |
| CIR1A_HUMAN | CIRH1A | 18 | 3 | 2,368 | 9,20E-03 | 0,0003113 | TRUE |
| WDR75_HUMAN | WDR75 | 19 | 3 | 2,333 | 1,42E-02 | 0,0003839 | TRUE |
| HNRL2_HUMAN | HNRNPUL2 | 10 | 1,5 | 2,33 | 0,002178 | 0,01902 | TRUE |
| K0020_HUMAN | KIAA0020 | 30,3 | 5,5 | 2,175 | 9,88E-05 | 8,55E-03 | TRUE |
| ILF3_HUMAN | ILF3 | 10 | 2 | 2,07 | 0,002661 | 0,02227 | TRUE |
| PABP4_HUMAN | PABPC4 | 9 | 2 | 1,999 | 0,004173 | 0,03137 | TRUE |

| | | | | | | | |
|---|---|---|---|---|---|---|---|
| IMP4_HUMAN | IMP4 | 9,7 | 2 | 1,975 | 0,004685 | 0,0349 | TRUE |
| MAK16_HUMAN | MAK16 | 10,3 | 2,5 | 1,945 | 0,001999 | 0,01783 | TRUE |
| RBM28_HUMAN | RBM28 | 47,3 | 10,5 | 1,873 | 1,69E-06 | 4,59E-04 | TRUE |
| UTP6_HUMAN | UTP6 | 13,7 | 3 | 1,872 | 0,001316 | 0,01291 | TRUE |
| RL5_HUMAN | RPL5 | 31,7 | 7,5 | 1,743 | 3,56E-03 | 0,0001727 | TRUE |
| RBM34_HUMAN | RBM34 | 28 | 7 | 1,719 | 1,10E-02 | 0,0003442 | TRUE |
| DHX9_HUMAN | DHX9 | 14,7 | 4 | 1,7 | 0,001052 | 0,0111 | TRUE |
| RRP5_HUMAN | PDCD11 | 34 | 8 | 1,651 | 8,09E-03 | 0,0002906 | TRUE |
| BRX1_HUMAN | BRIX1 | 28 | 8 | 1,638 | 1,02E-02 | 0,0003321 | TRUE |
| WDR46_HUMAN | WDR46 | 27 | 7,5 | 1,618 | 2,53E-02 | 0,0005875 | TRUE |
| RL7_HUMAN | RPL7 | 32,7 | 9,5 | 1,583 | 4,02E-06 | 0,0001733 | TRUE |
| DCA13_HUMAN | DCAF13 | 20 | 5,5 | 1,576 | 0,0004916 | 0,006142 | TRUE |
| WDR36_HUMAN | WDR36 | 24,3 | 6,5 | 1,532 | 0,0002527 | 0,003664 | TRUE |
| NOC4L_HUMAN | NOC4L | 18 | 5 | 1,516 | 0,001558 | 0,01488 | TRUE |
| DDX56_HUMAN | DDX56 | 22,3 | 7 | 1,498 | 0,0002267 | 0,003409 | TRUE |
| WDR43_HUMAN | WDR43 | 27,3 | 8 | 1,45 | 0,0001539 | 0,002551 | TRUE |
| HP1B3_HUMAN | HP1BP3 | 14,3 | 4,5 | 1,439 | 0,00493 | 0,03615 | TRUE |
| NOL6_HUMAN | NOL6 | 15,3 | 5 | 1,438 | 0,003095 | 0,02565 | TRUE |
| NKRF_HUMAN | NKRF | 17 | 5,5 | 1,359 | 0,003691 | 0,02828 | TRUE |
| KRR1_HUMAN | KRR1 | 19,7 | 6,5 | 1,333 | 0,002113 | 0,01865 | TRUE |
| RL18A_HUMAN | RPL18A | 18 | 6 | 1,327 | 0,00328 | 0,02637 | TRUE |
| RL3_HUMAN | RPL3 | 53 | 18,5 | 1,209 | 4,06E-03 | 0,0001733 | TRUE |
| UT14A_HUMAN | UTP14A | 29,7 | 11 | 1,195 | 0,0004597 | 0,00602 | TRUE |
| U3IP2_HUMAN | RRP9 | 21,3 | 8,5 | 1,185 | 0,00231 | 0,01995 | TRUE |
| RL28_HUMAN | RPL28 | 19 | 7 | 1,18 | 0,005844 | 0,04126 | TRUE |
| NOP2_HUMAN | NOP2 | 46 | 18,5 | 1,138 | 1,83E-02 | 0,0004791 | TRUE |
| UTP18_HUMAN | UTP18 | 28 | 11 | 1,113 | 0,001272 | 0,01275 | TRUE |
| DDX50_HUMAN | DDX50 | 25 | 10 | 1,064 | 0,003614 | 0,02795 | TRUE |
| RRS1_HUMAN | RRS1 | 38,7 | 16,5 | 1,052 | 0,0002236 | 0,003409 | TRUE |
| RL6_HUMAN | RPL6 | 35,3 | 15 | 0,9857 | 0,001118 | 0,01161 | FALSE |
| DDX21_HUMAN | DDX21 | 46,7 | 19 | 0,98 | 0,0002742 | 0,003839 | FALSE |
| CEBPZ_HUMAN | CEBPZ | 34,7 | 15,5 | 0,9599 | 0,00132 | 0,01291 | FALSE |
| KI67_HUMAN | MKI67 | 31 | 13,5 | 0,9415 | 0,003375 | 0,02687 | FALSE |
| EBP2_HUMAN | EBNA1BP2 | 31 | 13 | 0,9034 | 0,006125 | 0,04288 | FALSE |
| HNRPM_HUMAN | HNRNPM | 74,7 | 41,5 | 0,5466 | 0,005309 | 0,03815 | FALSE |

**Table S2, related to Figure 5. Putative H3K4ox readers. List of putative H3K4ox readers identified by mass spectrometry.** Gene symbol and MS score are shown. Prot nm: protein name; T: average of spectral counts in H3K4ox peptide; C: average of spectral counts in irrelevant peptide. LogFC: log-fold change. Adj p: adjusted p-value. DEP: Differential Enrichment analysis of Proteomic data (pvalue 0.05, 2 spectral counts, logFC 1); T: true, F: false. Two biological replicates of each condition were performed.

| Target gene | Direction | Sequence (5'-3') | Specie | Use |
|---|---|---|---|---|
| **CDC6** | Forward | TGGATGTTTGCAGGAGAGCTA | Human | mRNA-qPCR |
| | Reverse | GCTCCTTCTTGGCTCAAGGT | | |
| **CHAF1B** | Forward | CGGGTCCCTCCAGCATT | Human | mRNA-qPCR |
| | Reverse | TACACGGGCTCCTTGTTGTG | | |
| **CLSPN** | Forward | CTCAACAGGTGAAGACAGGCT | Human | mRNA-qPCR |
| | Reverse | CTTAGACGATTCCTTTGCCG | | |
| **DDB1** | Forward | CAAAAGGATAGCGCTGCC | Human | mRNA-qPCR |
| | Reverse | TGCATTACCAGAGAGCCGT | | |
| **H2A1A** | Forward | CGCCAAGTCTAAGTCTCGC | Human | mRNA-qPCR |
| | Reverse | TCCGCTCTGCATAGTTTCC | | |
| **H2A.Z.1** | Forward | CGGAATTCGAAATGGCTG | Human | mRNA-qPCR |
| | Reverse | TGTCGATGAATACGGCCC | | |
| **H2A.Z.2** | Forward | GAACATGGCTGGAGGCAAA | Human | mRNA-qPCR |
| | Reverse | CAAGTGTCTGTGGATGCGG | | |
| **GR 1** | Forward | GTCTCACTGTGTCCCCCAA | Human | ChIP-qPCR |
| | Reverse | ACCAGACTAGCCAACAAAGCA | | |
| **GR 3** | Forward | TCCGTTTCTTCTGGACGAAC | Human | ChIP-qPCR |
| | Reverse | CTCCAGCGCGAACTTTGTA | | |
| **GR 5** | Forward | GCCAGGCATGCTCTACTTT | Human | ChIP-qPCR |
| | Reverse | TATTAATCCAAGGCCGGG | | |
| **GR 6** | Forward | AGTGAATGTTTCATTGAGTGCTTA | Human | ChIP-qPCR |
| | Reverse | CTTTCAAATGGGTTCTTGTGA | | |
| **GR 9** | Forward | TCAGCCCCTGGAATAGCT | Human | ChIP-qPCR |
| | Reverse | TCCACCTGTACAGCCAGC | | |
| **GR 10** | Forward | GGCTTGTGAAACCAAGTCCA | Human | ChIP-qPCR |
| | Reverse | CCGAAGCTGGCAGATCAC | | |
| **GR 11** | Forward | GCACCCAACCAATATGTCTTC | Human | ChIP-qPCR |
| | Reverse | AGAGATACAACCAACACAGTGCA | | |
| **HPRT** | Forward | CTGGCGTCGTGATTAGTGAT | Human | mRNA-qPCR |
| | Reverse | GGCTACAATGTGATGGCCT | | |
| **LEF1** | Forward | CCCGTGAAGAGCAGGCTAAA | Human | mRNA-qPCR |
| | Reverse | TCGTTTTCCACCTGATGCAG | | |
| **LOXL2** | Forward | CCCCCTGGAGACTACCTGTT | Human | mRNA-qPCR |
| | Reverse | TTCGCTGAAGGAACCACCTA | | |
| **NFIA** | Forward | ATGTGAACGCAAGAAGCAG | Human | mRNA-qPCR |
| | Reverse | ATTCATCCTGGGTGAGACAG | | |

| | | | | |
|---|---|---|---|---|
| **Pumilio** | Forward | GACCAGCAGAATGAGATGGTTC | Human | mRNA-qPCR |
| | Reverse | CATAAGGATGTGTGGATAAGGCA | | |
| **RASD1** | Forward | CCACCGCAAGTTCTACTCCA | Human | mRNA-qPCR |
| | Reverse | GGATGAAAACGTCTCCTGTGAG | | |
| **RBX1** | Forward | CGACAGACCGTGTGTTTCC | Human | mRNA-qPCR |
| | Reverse | AGGGCTACTGCATTCCACTTT | | |
| **RHOB** | Forward | GTGTGTCTGTTCGACTCCCC | Human | mRNA-qPCR |
| | Reverse | AGGGATATCAAGCTCCCGC | | |
| **RING1B** | Forward | AGCACAATAATCAGCAAGCACT | Human | mRNA-qPCR |
| | Reverse | GCTCCACTACCATTTTCAATCTG | | |
| **RNA Pol II** | Forward | CTGAGTCCGGATGAACTGGT | Human | ChIP-qPCR |
| | Reverse | ACCCATAAGCAGCGAGAAAG | | |
| **RP1L1** | Forward | TAAGAACATGGACCCTCGCC | Human | mRNA-qPCR |
| | Reverse | CTGCAGCGAGTCCACCTTT | | |
| **RUVBL1** | Forward | GCCCTGGAGTCTTCTATCGC | Human | mRNA-qPCR |
| | Reverse | CACTCGGTCCAGAAGGTCA | | |
| **RUVBL2** | Forward | GATCATGGCCACCAACC | Human | mRNA-qPCR |
| | Reverse | CAGGTCTATGGGGATGCC | | |
| **TP53INP1** | Forward | CGTCTGGGTACCTGAACGA | Human | mRNA-qPCR |
| | Reverse | AGAAGAGTCATTGTACGTGGGC | | |

**Table S3, related to STAR Methods**. **Primers for ChIP- and mRNA-qPCR.** Primers used for mRNA and ChIP analysis. List of primer sequences used in this study, shown in 5' to 3' direction.
